# Structural basis for conformational equilibrium of the catalytic spliceosome

**DOI:** 10.1101/2020.09.22.306647

**Authors:** Max E. Wilkinson, Sebastian M. Fica, Wojciech P. Galej, Kiyoshi Nagai

## Abstract

The catalytic spliceosome exists in equilibrium between the branching (B*/ C) and exon ligation (C*/ P) conformations. Here we present the electron cryo-microscopy reconstruction of the *Saccharomyces cerevisiae* C complex at 2.8 Å resolution and identify a novel C-complex intermediate (C_i_) that elucidates the molecular basis for this equilibrium. In the C_i_ conformation, the exon-ligation factors Prp18 and Slu7 are already bound before ATP hydrolysis by Prp16, which destabilises the branching conformation. Biochemical assays suggest these pre-bound factors prime C complex for conversion to C* by Prp16. A complete model of the Prp19-complex (NTC) shows how the NTC pre-recruits the branching factors Yju2 and Isy1 before branching. Prp16 remodels Yju2 binding after branching, allowing Yju2 to remain associated with the C* and P spliceosomes and promote exon ligation. Our results explain how Prp16 action modulates dynamic binding of step-specific factors to alternatively stabilise the C or C* conformation and establish equilibrium of the catalytic spliceosome.

## Introduction

The spliceosome produces mRNA by excising introns from pre-mRNAs in two phosphoryl transfer reactions – branching and exon ligation. The spliceosome assembles *de novo* on each pre-mRNA through protein and RNA interactions that recognise conserved sequences at exon-intron junctions, called splice sites^1^ (**Fig. 1a**). The U6 snRNA recognises the 5′-splice site (5′-SS), while the U2 snRNA pairs with the intron around the branch adenosine (brA), forming the branch helix^1,2^. The 5′-exon is stabilised in the active site by pairing with loop I of the U5 snRNA (refs. 3,4). The U2 and U6 snRNAs fold into a triple helix conformation that constitutes the active site and allows U6 snRNA to position two catalytic metals (refs. 1,5,6). This active site is stabilised by binding of the Prp19-associated complex^1,7^ (NTC). Branching occurs in the B* complex when the 2′-hydroxyl of the brA attacks the 5′-SS. The resulting C complex is remodelled by the DEAH-box ATPase Prp16 into the C* open conformation (**Fig. 1b**). Docking of the 3′-splice site (3′-SS) at the active site (refs. 8,9) forms the closed C* complex, which then catalyses exon ligation, when the 3′-hydroxyl of the 5′-exon attacks the 3′-SS, resulting in mRNA formation and excision of the lariat-intron (**Fig. 1a,b**). Exon ligation forms the post-catalytic P complex (**Fig. 1b**), from which the ATPase Prp22 releases the mRNA.

**Figure 1.**
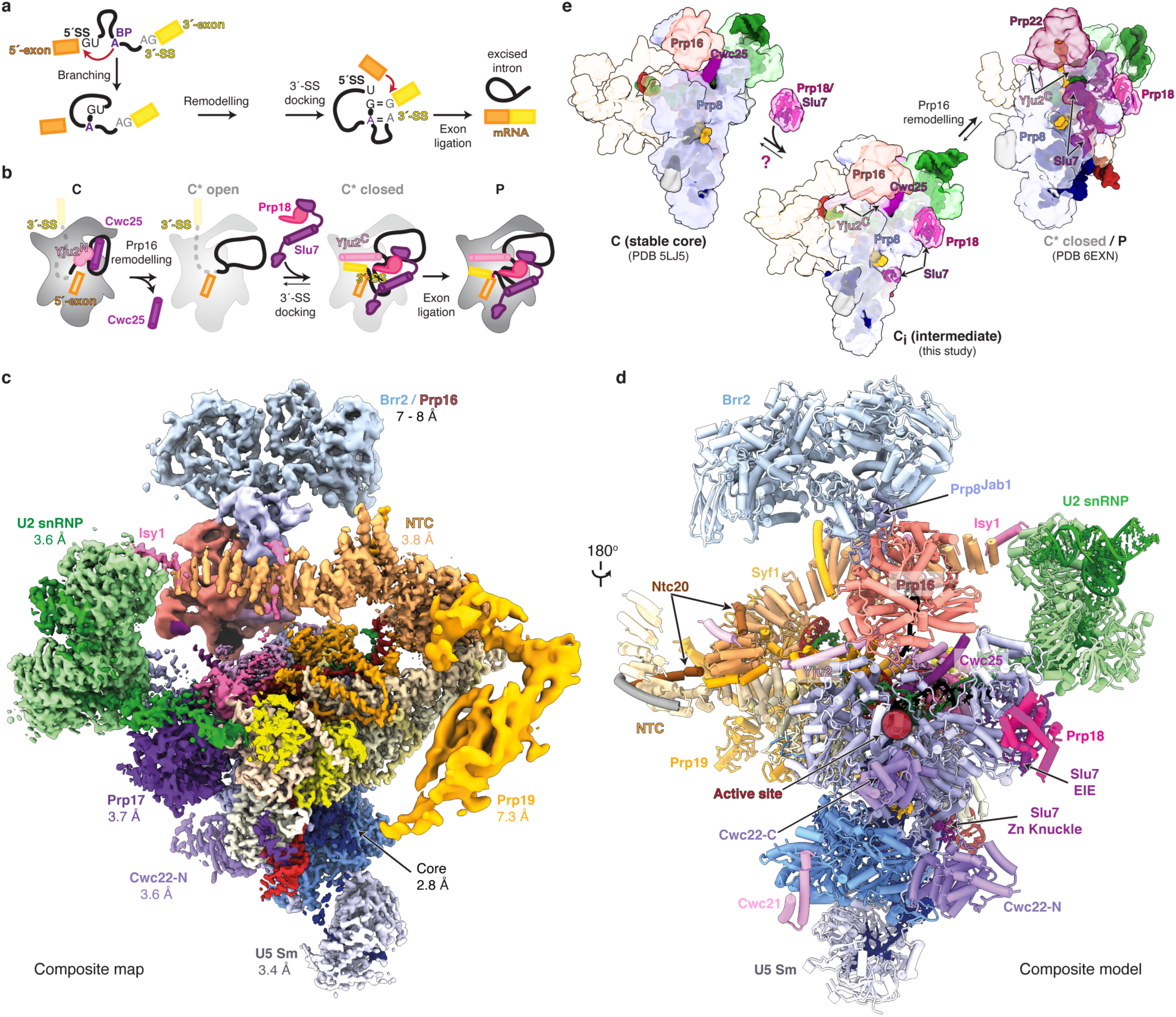
A novel C-complex spliceosome intermediate. **a**, Pre-mRNA splicing mechanism. **b**, Canonical pathway for binding of exon ligation factors during the catalytic stage of splicing. **c**, Composite cryo-EM map of the C-complex spliceosome. Average resolution (FSC 0.143) is indicated for each focussed map. **d**, Composite model of the C-complex spliceosome with pre-bound exon ligation factors. **e**, The novel C-complex intermediate (C_i_) suggests early binding of exon ligation factors before Prp16-mediated remodelling. The previously observed stable C conformation may exist in equilibrium with our novel C_i_ (intermediate) conformation.

Several step-specific splicing factors associate with the spliceosome during branching and exon ligation to promote splicing chemistry. In the budding yeast *Saccharomyces cerevisiae* the branching factors Cwc25, Yju2, and Isy1 (refs. 10-13) clamp the branch helix and dock it into the spliceosome active site^7,14,15^. The exon-ligation factors Prp18, Slu7 and Prp17, act more indirectly. Prp18 inserts a conserved loop near the active site to stabilise the docked 3′-SS, while Slu7 binds the spliceosome surface and rigidifies the C* conformation. Slu7 acts by an unknown mechanism to promote chemistry for substrates with a long distance between the brA and the 3′-SS (refs. 8,9,16-20). Prp17 binds the spliceosome before branching^21^ but functions primarily during exon ligation^22^ to stabilise an undocked conformation of the branch helix, as observed in the C* complex^8,9^.

While a single active site catalyses both splicing steps^5,7,9^, a structural rearrangement allows the 3′-SS to replace the branch helix during exon ligation. Genetic experiments have suggested a two-state model of the catalytic spliceosome where an equilibrium exists between the branching and exon - ligation conformations^23^. This equilibrium is modulated by Prp8 and by the ATPase Prp16 (refs. 24,25), whose activity promotes exon ligation in the forward direction by dissociating branching factors and undocking the branch helix from the active site (**Fig. 1b**, refs. 1,26). Indeed, under certain *in vitro* conditions, both catalytic steps of splicing are reversible, and disrupting the function of step-specific factors promotes reversal^27,28^. Intriguingly, the exon-ligation factors Slu7 and Prp18 have been suggested to bind to low affinity sites already in B*, while Prp16 action converts the binding sites to higher affinity sites in C* (ref. 29). Despite these biochemical and genetic studies, it remains unclear whether step-specific factors bind exclusively to either the branching or the exon-ligation conformation of the spliceosome.

We sought to determine the molecular basis for conformational equilibrium of the catalytic spliceosome by investigating whether branching and exon-ligation factors can engage the C-complex conformation at the same time. Here we present the detailed analysis of a large electron cryo-microscopy (cryo-EM) dataset of the C-complex spliceosome from budding yeast. The new cryo-EM reconstruction of the C complex at 2.8 Å resolution allowed building of the most complete atomic model of a catalytic spliceosome so far (**Fig. 1c,d**). The RNA-based active site is stabilised by both monovalent and divalent cations and uses non-Watson– Crick base pairs for splice site recognition. Most importantly, focussed classification revealed that exon-ligation factors Prp18 and Slu7 are pre-recruited to the branching conformation, creating a new C-complex intermediate (C_i_) (**Fig. 1e**), while biochemical assays suggest these pre-bound factors are competent to promote exon ligation upon conversion to the C* conformation by Prp16. Finally, a complete model of the NTC shows how Syf1 acts as a recruitment hub for the branching factors Yju2 and Isy1, whose binding is remodelled in the transition from branching to exon ligation. Thus, conformational equilibrium of the catalytic spliceosome is mediated by dynamic binding of step-specific factors to the C or C* conformation.

## Results

### The complete structure of the yeast C-complex spliceosome

We previously reported a 3.8 Å structure of the C-complex spliceosome stalled by a 3′-SS mutation and purified with affinity tags on Prp18 and Slu7 (ref. 7). While the 3′-SS UAc mutation was expected to stall spliceosomes mainly in the C* conformation^30,31^, accumulation of C complexes suggested that in the absence of stable 3′-SS docking the spliceosome may equilibrate back into the branching conformation (**Fig. 1b**) without complete dissociation of exon-ligation factors. Indeed, stalling the C* complex using a chemical modification of the 3′-SS also produced significant amounts of spliceosomes in the C conformation, which remained associated with Slu7 (ref. 8). These observations raised the possibility that exon-ligation factors may bind the C-complex spliceosome before Prp16 remodelling.

We merged all C-complex particles obtained by cryo-EM from various spliceosome purifications and investigated binding of Slu7 and Prp18 by focussed classification (**Extended Data Figs. 1-4**). This yielded a cryo-EM reconstruction at 2.8 Å resolution for the C-complex core, showing complete base separation, base and side chain rotamers, and allowing discrimination between adenosine and guanosine bases (**Extended Data Fig. 5**). This map allowed high-confidence modelling of every protein and RNA in the core of the catalytic spliceosome. The large number of particles facilitated classification and focussed refinement for the peripheral flexible regions of the spliceosome (**Extended Data Figs. 2, 3**). This produced improved maps for the U2 snRNP (3.6 Å resolution) and NTC (3.8 Å resolution) and allowed atomic modelling of these regions, which in all previous yeast spliceosome structures were only modelled by homology. The C-complex helicase module (Brr2 and Prp16) and Prp19 module (Prp19, Snt309, Cef1 C-terminus) were each refined to sub-nanometre resolution (6-7 Å), allowing discrimination of secondary structures and model building by molecular dynamics flexible fitting of crystal structures and homology models (**Extended Data Figs. 2-5, Table 1**). Together, these improved maps allow us to present the most complete molecular model of a catalytic spliceosome (**Fig. 1c,d**). The structure reveals novel features of the active site and unexpected binding of Prp18 and Slu7 to C complex, thus shedding light on remodelling during the catalysis.

**Table 1.** Cryo-EM data collection, refinement and validation statistics

|  | Core<br>(EMDB-xxxx) | U2 snRNP<br>(EMDB-xxxx) | NTC<br>(EMDB-xxxx) |
| --- | --- | --- | --- |
| **Data processing** |  |  |  |
| Particle images (no.) | 403,474 | 294,934 | 242,546 |
| Map resolution (FSC=0.143) (Å) | 2.80 | 3.58 | 3.82 |
| Map resolution range<sup>a</sup> (Å) | 2.71 – 4.94 | 3.39 – 8.20 | 3.64 – 8.68 |
| Map resolution after density modification (Å) | 2.69 | 3.26 | 3.50 |
| **Refinement** |  |  |  |
| Refined residues<sup>b</sup> | /6/E/I/C/J/K/N/M/P/L/F/a/c2:4-106/5:27-126/A:126-2078,2500/O:11-252/S:36-256/y:95-211/R:2-26/H:289-486/o:51-76/c:122-140,400/G:2-98/D:1-210 | /2:107-1109/Y/W/q/p/r/n/k/l/m/G:214-232 | /T/S:257-625/y:2-81/Z/G:163-191/K:1-11/J:46-59/X/O:304-327/D:209-225 |
| Model resolution (FSC=0.5) (Å) | 2.76 | 3.33 | 3.57 |
| Map CC (around atoms) | 0.68 | 0.75 | 0.69 |
| Map sharpening *B* factor (Å<sup>2</sup>) | -89 | -138 | -157 |
| Model composition |  |  |  |
| Non-hydrogen atoms | 51,668 | 9959 | 9030 |
| Protein residues | 5463 | 932 | 1260 |
| RNA residues | 342 | 119 | 0 |
| Ligands | 17 | 0 | 0 |
| Mean *B* factors (Å<sup>2</sup>) |  |  |  |
| Protein | 100.3 | 59.0 | 94.2 |
| RNA | 129.7 | 182.9 |  |
| Ligand | 104.8 |  |  |
| R.m.s. deviations |  |  |  |
| Bond lengths (Å) | 0.007 | 0.006 | 0.011 |
| Bond angles (°) | 1.203 | 1.195 | 1.339 |
| Validation |  |  |  |
| MolProbity score | 0.60 | 1.14 | 0.85 |
| Clashscore | 0.26 | 0.58 | 0.48 |
| Rotamer outliers (%) | 0.14 | 1.77 | 0.27 |
| CaBLAM outliers (%) | 0.78 | 0.91 | 0.87 |
| RNA geometry |  |  |  |
| Correct sugar puckers (%) | 97.1 | 96.6 |  |
| Good backbone conformations (%) | 79.8 | 79.8 |  |
| Ramachandran plot |  |  |  |
| Favoured (%) | 98.56 | 96.25 | 96.94 |
| Allowed (%) | 1.42 | 3.42 | 3.06 |
| Disallowed (%) | 0.02 | 0.33 | 0.00 |

<sup>a</sup> Range from 5<sup>th</sup> to 95<sup>th</sup> percentiles of local resolution map within the refinement mask
<sup>b</sup> Shown in ChimeraX selection notation, where / indicates chain ID and : indicates residue IDs, e.g. /2:4-47 is chain 2 (U2 snRNA) residues 4 – 47.

**Table 1 (cont.) | Cryo-EM data collection, refinement and validation statistics**
|  | U5 Sm<br>(EMDB-xxxx) | Cwc22 NTD<br>(EMDB-xxxx) | Helicase module<br>(EMDB-xxxx) |
| --- | --- | --- | --- |
| **Data processing** |  |  |  |
| Particle images (no.) | 65,561 | 73,802 | 103,461 |
| Map resolution (FSC=0.143) (Å) | 3.13 overall<br>3.4 around U5 Sm | 3.16 overall<br>3.6 around<br>Cwc22N | 7.15, 8.0<br>(composite map) |
| Map resolution range<sup>a</sup> (Å) | 3.06 – 8.41<br>around U5 Sm | 3.40 – 5.24<br>around Cwc22N | 6.43 – 10.56 |
| Map resolution after density<br>modification (Å) | 2.96 | 2.97 |  |
| **Refinement** |  |  |  |
| Refined residues<sup>b</sup> | /5:1-26,127-178/b/d/e/f/g/h/j | /H:10-264/X:1-20 | /A:2150-2395/B/Q/I:87-92 |
| Model resolution (FSC=0.5) (Å) | 4.18 | 4.29 | 8.57 |
| Map CC (around atoms) | 0.60 | 0.64 | 0.65 |
| Map sharpening *B* factor (Å<sup>2</sup>) | -74 | -89 | -275, -494 |
| Model composition |  |  |  |
| Non-hydrogen atoms | 6424 | 2166 | 20,741 |
| Protein residues | 602 | 273 | 2577 |
| RNA residues | 78 | 0 | 6 |
| Ligands | 0 | 0 | 0 |
| Mean *B* factors (Å<sup>2</sup>) |  |  |  |
| Protein | 162.9 | 154.39 | Not refined |
| RNA | 257.7 |  | Not refined |
| Ligand |  |  |  |
| R.m.s. deviations |  |  |  |
| Bond lengths (Å) | 0.006 | 0.006 | 0.012 |
| Bond angles (°) | 1.197 | 1.171 | 1.945 |
| Validation |  |  |  |
| MolProbity score | 0.99 | 0.67 | 1.09 |
| Clashscore | 0.49 | 0.23 | 1.13 |
| Rotamer outliers (%) | 1.85 | 0.00 | 0.82 |
| CaBLAM outliers (%) | 0.88 | 0.40 | 1.61 |
| RNA geometry |  |  |  |
| Correct sugar puckers (%) | 93.6 |  | 83.3 |
| Good backbone conformations (%) | 71.8 |  | 83.3 |
| Ramachandran plot |  |  |  |
| Favoured (%) | 97.43 | 97.62 | 96.06 |
| Allowed (%) | 2.40 | 2.38 | 3.74 |
| Disallowed (%) | 0.17 | 0.00 | 0.19 |

<sup>a</sup> Range from 5<sup>th</sup> to 95<sup>th</sup> percentiles of local resolution map within the refinement mask
<sup>b</sup> Shown in ChimeraX selection notation, where / indicates chain ID and : indicates residue IDs, e.g. /2:4-47 is chain 2 (U2 snRNA) residues 4 – 47.

**Table 1 (cont.) | Cryo-EM data collection, refinement and validation statistics**
|  | Prp17 WD40<br>(EMDB-xxxx) | Prp19<br>(EMDB-xxxx) | Combined<br>model<br>(EMDB-xxxx)<br>(PDB-NNNN) |
| --- | --- | --- | --- |
| **Data processing** |  |  |  |
| Particle images (no.) | 59,152 | 49,514 | NA |
| Map resolution (FSC=0.143) (Å) | 3.67 overall | 7.30 | Composite map |
| Map resolution range<sup>a</sup> (Å) | 3.27 – 4.74<br>around Prp17 | 6.79 – 9.48 | 2.85 – 10.6 |
| **Refinement** |  |  |  |
| Refined residues<sup>b</sup> | /o:149-454 | /O:484-536/s/t/u/v/w/X:35-51 | All |
| Model resolution (FSC=0.5) (Å) | Not determined | Not determined | 3.1 |
| Map CC (around atoms) | 0.43 | 0.39 | Not determined |
| Map sharpening *B* factor (Å<sup>2</sup>) | -90 | -200 | - |
| Model composition |  |  |  |
| Non-hydrogen atoms | 2454 | 6729 | 109,171 |
| Protein residues | 304 | 906 | 12,317 |
| RNA residues | 0 | 0 | 545 |
| Ligands | 0 | 0 | 17 |
| Mean *B* factors (Å<sup>2</sup>) |  |  |  |
| Protein | Not refined | Not refined | 134.12 |
| RNA |  |  | 156.50 |
| Ligand |  |  | 104.8 |
| R.m.s. deviations |  |  |  |
| Bond lengths (Å) | 0.013 | 0.023 | 0.010 |
| Bond angles (°) | 2.180 | 2.566 | 1.513 |
| Validation |  |  |  |
| MolProbity score | 0.95 | 1.09 | 0.82 |
| Clashscore | 0.21 | 2.36 | 0.65 |
| Rotamer outliers (%) | 1.09 | 0.90 | 0.59 |
| CaBLAM outliers (%) | 2.70 | 0.48 | 1.00 |
| RNA geometry |  |  |  |
| Correct sugar puckers (%) |  |  | 96.3 |
| Good backbone conformations (%) |  |  | 78.3 |
| Ramachandran plot |  |  |  |
| Favoured (%) | 95.00 | 97.67 | 97.47 |
| Allowed (%) | 5.00 | 2.33 | 2.45 |
| Disallowed (%) | 0.00 | 0.00 | 0.08 |

<sup>a</sup> Range from 5<sup>th</sup> to 95<sup>th</sup> percentiles of local resolution map within the refinement mask
<sup>b</sup> Shown in ChimeraX selection notation, where / indicates chain ID and : indicates residue IDs, e.g. /2:4-47 is chain 2 (U2 snRNA) residues 4 – 47.

### Monovalent metal ion binding in the spliceosome active site

The improved resolution of our new C-complex map allows a finer analysis of the active site of the spliceosome. The active site is formed by the U6 snRNA, which adopts a triplex conformation to position two catalytic Mg^2+^ ions during branching and exon ligation^1,5,6,32^. The spliceosome uses the same configuration of the active site and the same two-metal ion mechanism for catalysis as observed in group II self-splicing introns^5,33^, and recent structural studies support the evolution of the spliceosome from group II introns^34,35^. The active site of group II introns binds a composite, trinuclear metal cluster, in which the classical two catalytic Mg^2+^ ions (M1 and M2) are stabilised by a third monovalent ion, usually K^+^ (K1) (**Extended Data Fig. 6**, refs. 33,36), which is necessary for efficient branching by some group II introns^33^. Thus, an additional positive charge may be required to stabilise M1 and M2 binding during catalysis. Although several spliceosome structures were reported with non-catalytic Mg^2+^ ions stabilising the U6 snRNA triplex conformation^15,37^, no unambiguous density for K1 had been observed so far, despite several studies suggesting pre-mRNA splicing requires monovalent cations^38,39^.

In our 2.8 Å map of C complex, we observed a new spherical density clearly separated from adjacent phosphates and bases in a position corresponding to K1 in group II introns (**Fig. 2a-c, Extended Data Fig. 6**). This new density has octahedral coordination geometry from surrounding oxygen ligands, with bond distances of 2.7 to 3 Å, consistent with a monovalent cation^40^ (**Fig. 2c**). As our spliceosomes were purified using buffers containing K^+^, we assigned this density as a potassium ion. This ion interacts with three of the four U6 snRNA residues that provide ligands for the catalytic Mg^2+^ ions, including all the ligands for M2 (A59, G60, and U80), as well as with the third strand of the triplex (G52) and is thus ideally positioned to stabilise M1 and M2 binding at the active site, as does K1 in group II introns (**Fig. 2a,b**). Intriguingly, right after formation of the active site triplex, in B^act^, this new K1 site is partially blocked by a lysine from Prp11, a U2 snRNP SF3 sub-complex protein that is dissociated by Prp2 during catalytic activation^37^ (**Extended Data Fig. 6**). In the late B* complex, before productive docking of brA at the active site, the K1 site remains unoccupied^15^. The C complex is the first spliceosome state where K1 is fully occupied and this site remains filled in P complex cryo-EM maps, provided that K^+^ is present in the purification buffers^18^ (**Extended Data Fig. 6**). Such progressive occupancy of the K1 site appears conserved in human spliceosomes (**Extended Data Fig. 6**) and may couple catalytic competence with docking of the branch helix at the active site. We propose that formation of the full M1-M2-K1 metal cluster is linked to docking of the brA at the active site and to catalytic activation of the spliceosome. This metal cluster is maintained throughout the catalytic stage and promotes both branching and exon ligation, similarly to its group II intron equivalent^33^.

**Figure 2.**
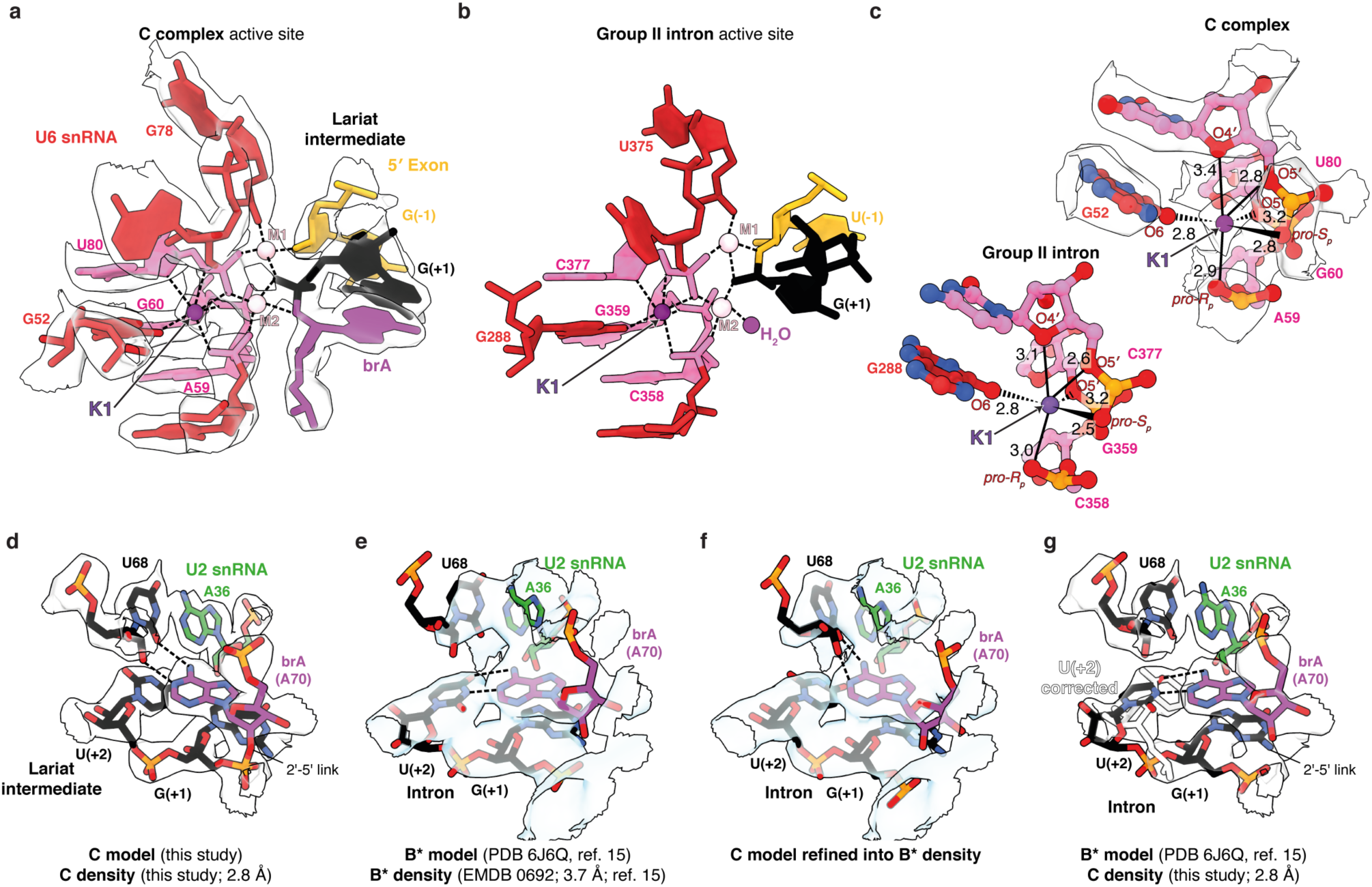
Key active site elements of the C complex. **a**, A potassium ion stabilises the catalytic centre of the spliceosome. Metal coordination is shown as dashed lines; cryo-EM density is shown as transparent contour surface. **b**, Structure of the M1-M2-K1 catalytic metal cluster in a group IIa intron before hydrolysis (PDB 4FAQ, ref. 33). **c**, Details of the K1 coordination sphere in the spliceosome and group II intron. Distances are labelled in Ångstrom. **d-g**, Recognition of the branch adenosine (brA) during branching. Comparison of our proposed brA pairing in B* and C with the previously suggested brA interactions in B*.

### Non-Watson–Crick base pairs recognise the branch adenosine during branching

The brA bulges out of the branch helix formed between the intron and U2 snRNA and must be precisely positioned to allow for nucleophilic attack on the 5′-SS (refs. 1,7). Thus, recognition of the brA is essential for efficient branching, and specific mutations of this adenosine inhibit branching *in vitro* and *in vivo*^41,42^ during the catalytic stage^25^ by destabilising the branching conformation^23^. In our original 3.8 Å structure of C complex^7^, we built the branch adenosine such that its Watson–Crick face paired with the sugar edge of intron nucleotide U(+68), two nucleotides upstream (UAC<u>U</u>A**A**C, brA in bold, U(+68) underlined). However, in a different, 3.4 Å structure of C complex^14^ this pairing was not modelled. Instead, in subsequent structures of a B* complex poised for branching^15^ a canonical Watson–Crick base pair was proposed between brA and the 5′-SS nucleotide U(+2), although in these structures the local resolution around the brA was limited (∼ 4 Å) (**Fig. 2e**). Therefore, the structural basis for the nucleotide specificity of the branching reaction was unclear.

In our new structure of C complex, the branch helix is robustly docked by the branching factors and is visible at a local resolution of 2.8 Å around the branch adenosine. The density unambiguously demonstrates that brA pairs to the sugar edge of U(+68) in a base-triple interaction, while the 5′-SS U(+2) pairs with U2 snRNA G37 in the context of another base triple (**Fig. 2c**), as we proposed in our earlier model^7^. Furthermore, this model can be convincingly refined into density for the B* complex poised for branching, with Yju2 and Isy1 bound to the branch helix^15^ (**Fig. 2d,e**). Indeed, mutations at U(+2) of the intron primarily impair exon ligation, which is inconsistent with a functional role for U(+2) in pairing to brA during branching (refs. 23,24). The previous B* model likely originates from the ambiguity of map interpretation at approximately 4 Å resolution and there is no evidence for brA base-pair remodelling between B* and C complexes (**Fig. 2e-g**). Here, we propose that the brA is recognised for branching through non-canonical pairing with U(+68), which anchors it to the branch helix while allowing the 2′-hydroxyl to be positioned for nucleophilic attack on the 5′-SS. Our model explains the effects of specific branch adenosine mutations. Substitution of brA with cytosine or uridine strongly inhibits branching, while mutation to guanosine has milder effects *in vivo* and *in vitro*^23,25,43^, and these effects correlate with the level of distortion and movement of the brA 2′-OH that would result from accommodating base pairs analogous to the interaction of brA to U(+68) (ref. 44, **Extended Data Fig. 7**). These observations further underscore the importance of careful analysis and interpretation of cryo-EM maps in the context of previous functional studies of the spliceosome^45^.

### The exon-ligation factors Prp18 and Slu7 are recruited to a new on-pathway C_i_ intermediate

In C* and P complexes, the exon-ligation factor Prp18 binds to a face of the Prp8 RNaseH domain (Prp8^RH^) that would also be accessible in C complexes (**Figs. 1b, 3a**), suggesting that Prp18 might bind the spliceosome even before Prp16-mediated remodelling. We wondered whether such early binding of Prp18/ Slu7 explained how we could purify C complexes using affinity tags on Prp18 or Slu7 (refs. 7,8). Indeed, by focussed classification we identified a subset of particles in the C-complex conformation with extra density on the same face of Prp8^RH^ that Prp18 binds in C* (**Figs. 1d, 3a, c, Extended Data Figs. 3, 8**). The alpha-helical domain of Prp18 visible in C* was a perfect fit for this additional density, which also accommodated a small peptide of Slu7 (termed EIE element, **Fig. 3b,c**) that is known to bind Prp18 (ref. 19). The conserved loop of Prp18 that stabilises the 3′-SS docked in the active site in C*/P is not visible in this subset of particles (Prp18^CR^, ref. 9, **Fig. 3c,d**), suggesting it is disordered before Prp16 remodelling and 3′-SS docking. Biochemical experiments indicate that endogenous Prp18 and Slu7 may exist as a heterodimer^19,46^ and thus early binding of Prp18 would also imply early binding of Slu7. Indeed, we also observed density corresponding to the Zn knuckle of Slu7 (ZnK) bound to Prp8 in this new intermediate (**Figs. 3b,c Extended Data Fig. 8**). This novel intermediate between C and C*, which we refer to as C_i_, demonstrates that all branching and exon-ligation factors can be bound in one complex, in agreement with previous biochemical data^29^.

**Figure 3.**
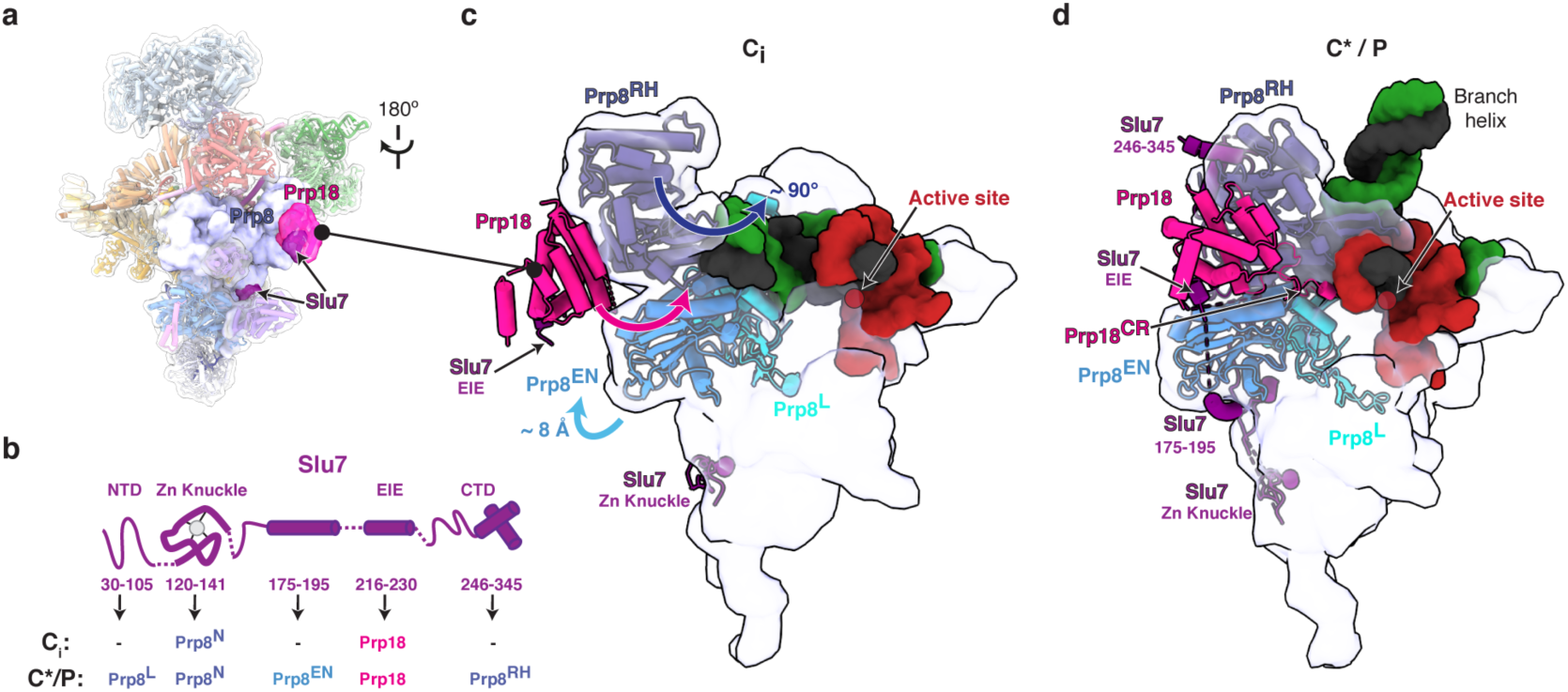
Prp18 and Slu7 are recruited to a new C_i_ conformation. **a**, Location of exon-ligation factors in the C-complex spliceosome. Prp8 and Prp18/ Slu7 are shown as surface representation. **b**, Domain architecture of Slu7 and the interaction partners of each domain. **c-d**, Pre-recruited Prp18 and Slu7 are remodelled from the C_i_ to the C*/P conformation. Upward movement of the Prp8 EN domain creates additional binding sites for Slu7 in C*/P. The Prp18 conserved region (CR) is disordered in C_i_ but engages the active site in C*/P.

To address the functional relevance of early binding of Prp18/ Slu7 before Prp16 remodelling, we investigated whether the C_i_ intermediate could be purified independently of Prp18 and Slu7, using a branching factor that binds stably to the C complex before Prp16 action. We assembled native complexes on wild-type pre-mRNA and stalled the C complex with a cold-sensitive Prp16 mutation (*prp16-302*) that impairs ATP hydrolysis (C^*prp16-302*^, **Fig. 4a**). We then purified the stalled C-complex spliceosomes with the branching factor Cwc25, which remains bound after branching when Prp16 action is blocked^12,27^. Cryo-EM reconstruction revealed that these C spliceosomes were structurally identical to the C complex assembled on the 3′-SS UAc mutant substrate (**Fig. 4b, Extended Data Figs. 2, 8**). Moreover, focussed classification identified a subset of particles in the C_i_ conformation, in which Cwc25 and Prp18/ Slu7 were bound to the same complex (**Fig. 4c, Extended Data Figs. 3, 8**). Therefore, Prp18 and Slu7 can bind to the spliceosome before Prp16 action and the C_i_ complex can be purified with a branching factor.

**Figure 4.**
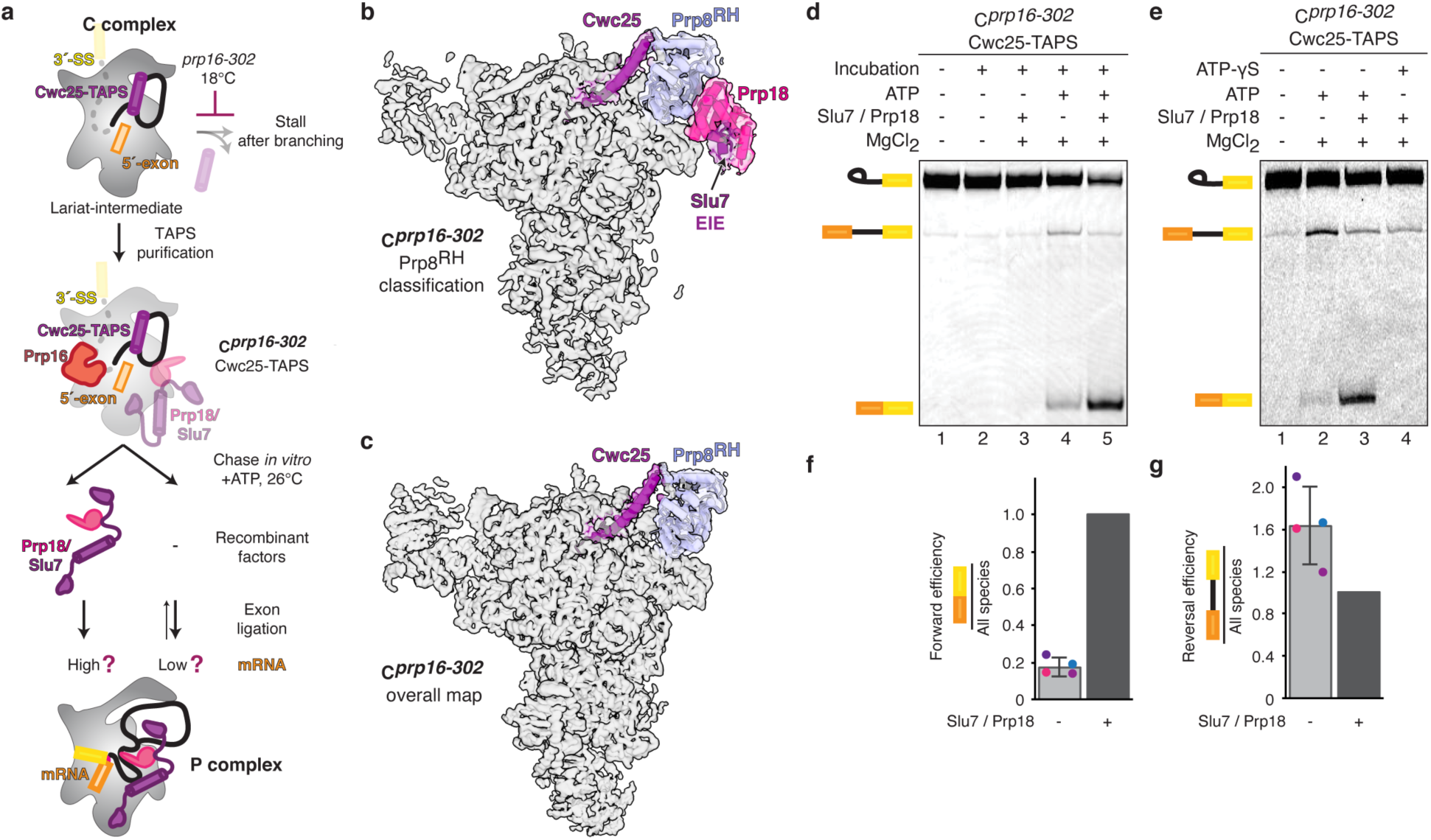
The novel C_i_ conformation is an on-pathway intermediate. **a**, Biochemical assay to assess function of the C_i_ conformation. Note that *prp16-302* impairs ATP hydrolysis at low temperatures and stalls Prp16 on the spliceosome right after branching. Chase of the purified C^*prp16-302*^ complex at the permissive temperature is predicted to allow mRNA production even in the absence of recombinant exon ligation factors. **b-c**, Focussed classification reveals a subpopulation of C^*prp16-302*^ spliceosomes in the C_i_ conformation. Maps were reconstructed without refinement from the classified particles and filtered to 5Å to aid comparison of additional density for Prp18/ Slu7. **d**, Purified C^*prp16-302*^ complexes produce mRNA when chased in the presence of ATP; the substrate was 3′-end labelled with Cy2 prior to spliceosome assembly. **e**, ATP hydrolysis is required for both exon ligation and branching reversal by C^*prp16-302*^ spliceosomes. **f-g**, Quantification of exon ligation and branching reversal efficiency for C^*prp16-302*^ spliceosomes. Relative ratios were normalised in each experiment to the condition where recombinant Slu7/ Prp18 was added. Values are averages of four independent experiments from three independent purifications; error bars represent s.d. (n=4). Values for individual experiments are indicated as dots overlaid on the bar graphs.

Overall, an average of 30% of C complexes resulting from various stalls during catalysis are in a C_i_ conformation and contain Prp18 and Slu7. However, only the small EIE element of Slu7 that tightly interacts with Prp18 is observed in all C_i_ particles while the ZnK of Slu7 engages only a subpopulation of C_i_ complexes (**Fig. 1b, Extended Data Fig. 8**). Indeed, binding of Prp18 and of the Slu7 ZnK can occur independently of each other, but focussed classification indicated that binding of Prp18 significantly increases the odds for binding of the Slu7 ZnK, and vice versa (odds ratio = 1.35). This relationship was observed regardless of the stall used to obtain the C_i_ intermediate, providing evidence that Prp18 and Slu7 interact as a heterodimer independently of their binding to the spliceosome (**Extended Data Fig. 8**). This mode of binding may allow early recruitment of Prp18 and/or Slu7 while ensuring that stable engagement of both exon-ligation factors with the spliceosome core does not occur until after Prp16 remodelling.

To determine whether C_i_ is a functional on-pathway intermediate, we assayed whether C^*prp16-302*^ complexes could produce mRNA when chased at the permissive temperature, as the *prp16-302* mutation allows growth and therefore splicing *in vivo* at 25°C (ref. 47). As expected, incubation of the purified C^*prp16-302*^ complexes at 25°C without ATP or with non-hydrolysable ATP-γS did not allow exon ligation, demonstrating that these complexes are stalled before Prp16 has acted to allow remodelling to C* (**Fig. 4d,e**, ref. 30). Incubation with ATP and exogenous, recombinant Prp18/ Slu7 resulted in efficient exon ligation, showing the spliceosomes are on-pathway intermediates. Importantly, incubation with ATP in the absence of exogenous Prp18/ Slu7 still produced significant levels of mRNA (**Fig. 4d-f**), demonstrating that the small C_i_ population containing Prp18/ Slu7 (**Extended Data Fig. 8**) stalled by the *prp16-302* mutation represents a functional on-pathway intermediate. Since Prp18/ Slu7 are necessary for UBC4 splicing^26^, we infer that the Prp18/ Slu7 pre-bound in C_i_ are competent to promote exon ligation upon remodelling by Prp16. Intriguingly, in the absence of exogenous Prp18/ Slu7, C^*prp16-302*^ complexes also catalysed reverse branching producing pre-mRNA (**Fig. 4d-g**). Thus, when Prp18 and Slu7 are limiting, the C_i_ spliceosome can revert to the B* conformation, supporting the idea that Prp18 and Slu7 shift the equilibrium towards the C* conformation during remodelling^8,9,26^.

### Complete structure of the Prp19-associated complex (NTC) reveals basis for recruitment of branching factors

The NTC is one of the biggest pre-assembled components of the spliceosome and joins during formation of the active site triplex in the B^act^ complex^1,48^. The NTC stabilises the U6 triplex^6^ and is essential for correct docking of the 5′-SS in the active site^49,50^. Moreover, branching factors were initially identified as proteins that interact loosely with the NTC (refs. 10,11,13,51,52). Previous yeast spliceosome structures revealed the overall architecture of the NTC as a sprawling clamp centred on a hinge formed by the tetrameric helical bundle of Prp19, from which Syf1 and Clf1 emerge as two helical arches that allow other components such as Cef1 to engage the spliceosome core (**Fig. 5a**). The Syf1 and Clf1 arches are very mobile during the C to C* transition^1,48^ and this feature has reduced local resolution for these regions in previous spliceosome maps. Thus, most of Clf1 and Syf1 were previously built only as idealised alpha helices of uncertain register.

**Figure 5.**
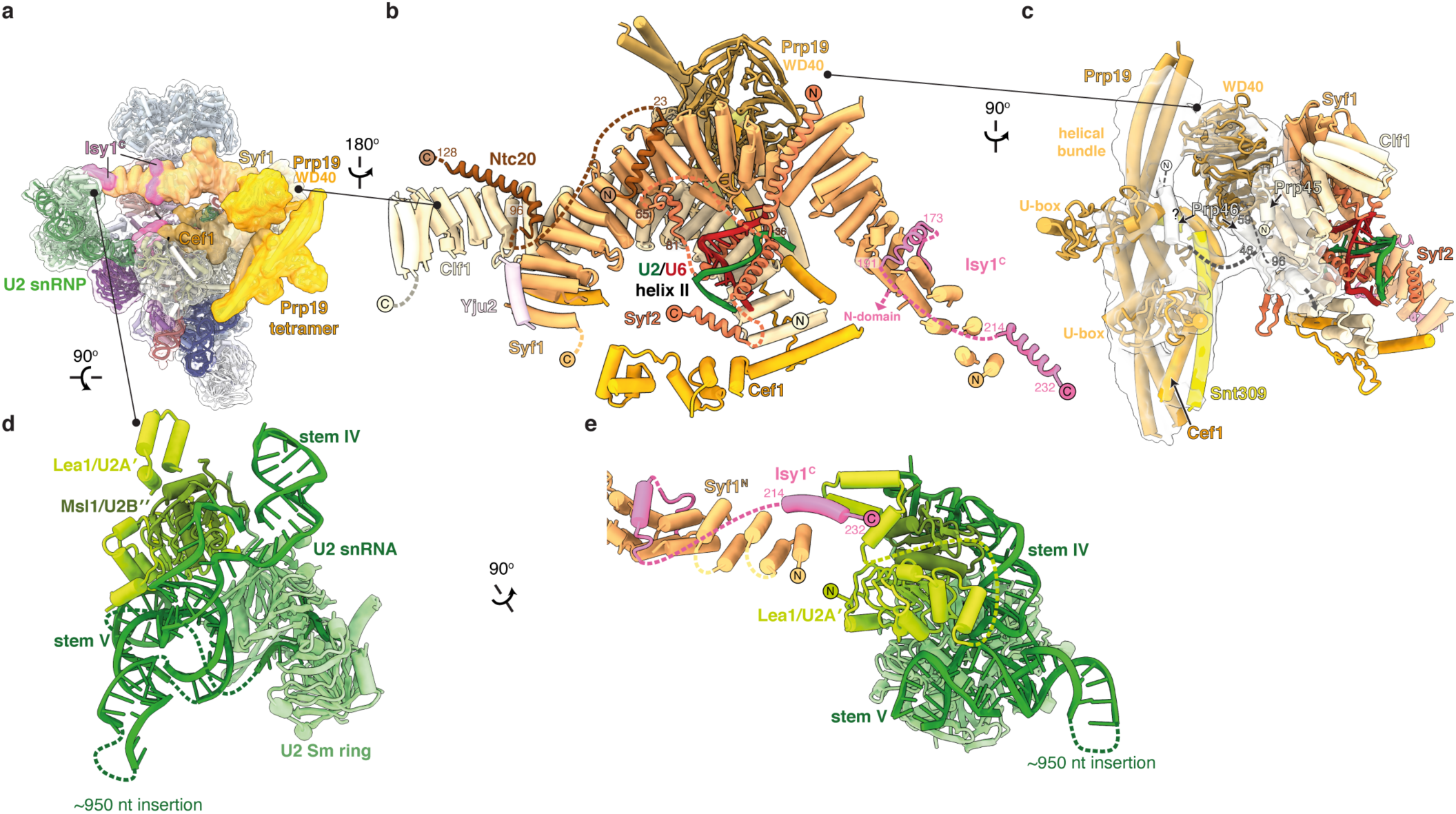
Complete architecture of the NTC and U2 snRNP in C complex. **a**, Location of NTC and U2snRNP in C complex. **b**, Architecture of the NTC showing newly identified Ntc20, Syf2, and Isy1 elements. **c**, The N-termini of Prp45 and Prp46 mediate attachment of the Prp19 helical bundle to the spliceosome. Focus-classified Prp19 density is shown as transparent contour surface. **d**, Complete structure of the U2 snRNP in C complex. Note that yeast contain a large, flexible insertion in U2 snRNA that is not visible in our map. **e**, Isy1 bridges the NTC to U2 snRNP through Lea1.

Our large C-complex dataset allowed focussed refinement that significantly improved density quality for Syf1 and Clf1 (**Extended Data Fig. 3**), showing surprisingly that several of the alpha helices previously assigned to Syf1 come from NTC proteins Syf2, Isy1, and Ntc20, and the U2 snRNP protein Lea1. We were able to build the N-terminal domain of Syf2 for the first time, revealing how Syf2 bridges Syf1 and U2/U6 helix II to anchor the NTC to active site elements in the core through Cef1 (**Fig. 5b**). Three helices on the surface of Clf1 were assigned to the N-termini of Prp45 and Prp46, with the latter projecting towards the Prp19 helical bundle, possibly acting as a tether for this domain (**Fig. 5c**). Ntc20 was the only unassigned NTC protein and here we show that two helices from Ntc20 bind the C-terminal domains of Syf1 and Clf1, respectively (**Fig. 5b, Extended Data Fig. 5**), thus linking these arches during activation and catalysis, consistent with previous genetics and biochemistry experiments^51,52^.

Our new map also explains how Isy1 (also known as Ntc30) is recruited to the spliceosome to act as a branching factor. The N-terminal domain of Isy1 was previously built in C complex as a clamp element that promotes docking of the branch helix at the active site^7,14^. We identified two helices from the C-terminus of Isy1 that bind the N-terminus of Syf1, explaining previous biochemical and genetic data implicating Isy1 as a peripheral NTC component^51^. Thus, the Isy1 N-terminal domain, which promotes branching, is likely already bound in B^act^ and tethered close to its site of action after Prp2 activity, in a manner parallel to Prp18 binding before exon ligation in C_i_. Our improved model of the U2 snRNP (**Fig. 5a,d**) shows that the C-terminal helix of Isy1 also bridges the extreme N-terminus of Syf1 to a newly identified 3-helix bundle of the U2 snRNP component Lea1 (**Fig. 5d,e**). Thus, Isy1 acts as a link between the NTC and the U2 snRNP and this interaction may be maintained throughout the catalytic stage and may influence C to C* remodelling, although Isy1 has yet to be identified in maps of C* or P complexes.

## Discussion

Our high-resolution structure of the C complex shows how the single active site of the spliceosome positions a trinuclear Mg^2+^ / K^+^ cluster for catalysis (**Fig. 2**). The active site remains catalytically licensed from B* onwards (**Extended Data Fig. 6**) but juxtaposes different reactants for each catalytic step. Although the 5′-exon remains bound onto U5 snRNA for both steps, the branch helix and brA must be removed from the active site after branching to make space for the 3′-SS to dock near the 5′-exon during exon ligation^1^. The ATPase Prp16 drives remodelling of brA interactions after branching to allow brA pairing to the 3′-SS, while Prp8 cradles the active site and governs a dynamic equilibrium between the branching and exon ligation conformations^2,23,24^. Step-specific factors also modulate this equilibrium^1,12,27-29^. The branching factors Cwc25 and Isy1 dissociate from the C-complex active site to allow exchange of reactants in the active site in C*/ P (ref. 1), while the exon-ligation factors Slu7 and Prp18 stabilise the C*/ P conformation to promote docking of the 3′-SS (ref. 1). Unexpectedly, we found that approximately 30% of C complex particles that retained branching factors (e.g. Cwc25) also contained the exon-ligation factor Prp18 (**Extended Data Fig. 8**), demonstrating that exon-ligation factors can bind the branching conformation before Prp16 remodelling. This new C_i_ intermediate, in which branching and exon-ligation factors bind the same complex (**Figs. 1e, 6**), provides the structural basis for how the spliceosome toggles between the branching and exon-ligation conformations.

In C_i_, Prp18 binds the Prp8^RH^ domain before Prp16 remodelling, and potentially before branching catalysis^29^. Slu7 is recruited to C_i_ through interactions of its EIE element with Prp18 or through binding of its zinc knuckle domain to the N-terminal domain of Prp8 (**Fig. 6**). Binding of the remainder of Slu7 is prevented in C_i_ by Cwc25, which would clash with the C-terminal domain of Slu7, while engagement of the central region of Slu7 requires upward movement of the Prp8^EN^ domain in C* (**Fig. 3**). Our C_i_ structure explains why previous studies have observed binding of Slu7 and Prp18 before Prp16 action at low salt concentrations and thus proposed a weaker affinity site before remodelling by Prp16 (ref. 29). In fact, our data suggest that exon-ligation factors bind at the same sites before and after Prp16 action but their affinity likely increases following remodelling by Prp16 (**Fig. 6**).

**Figure 6.**
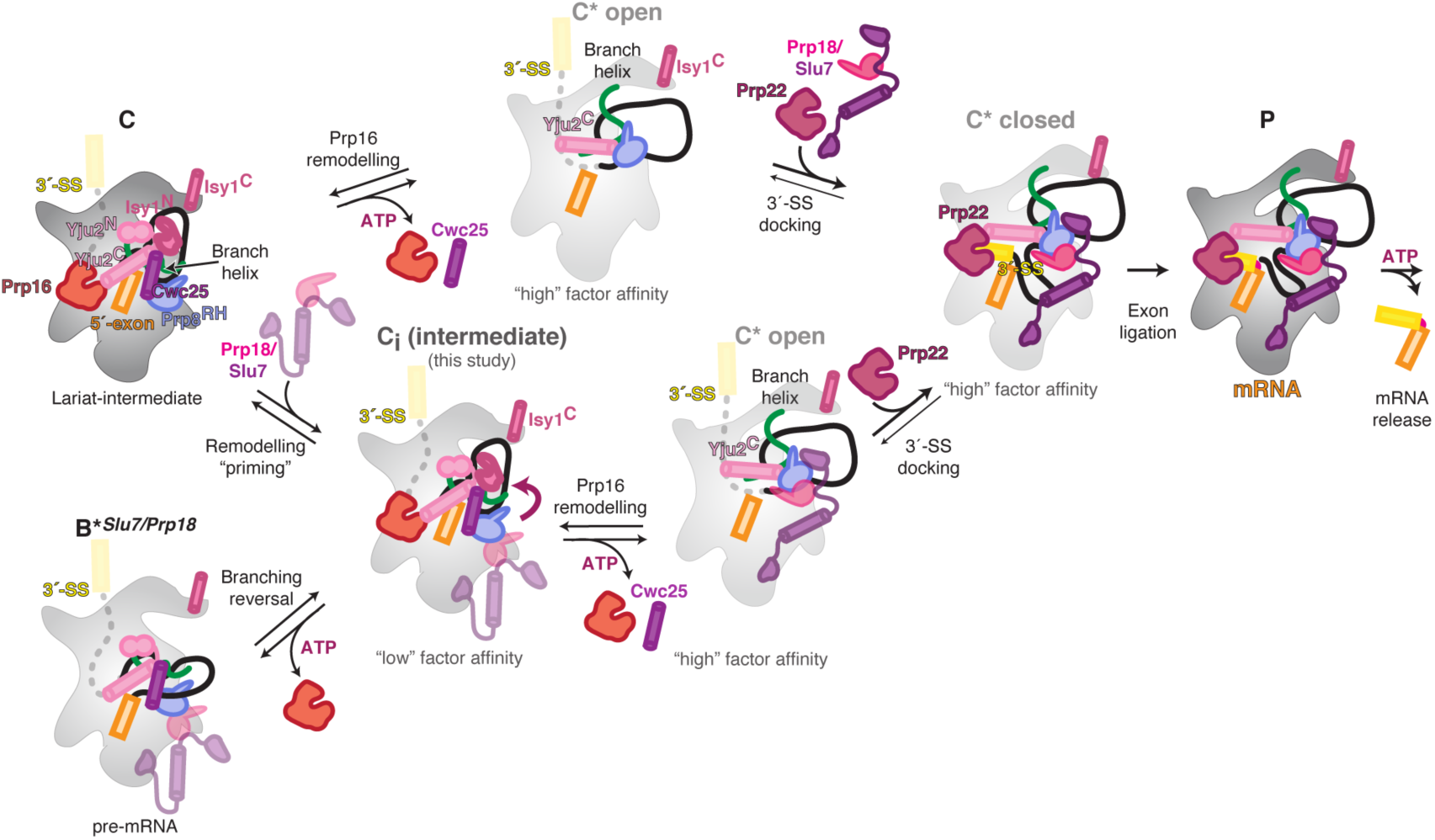
Model for conformational equilibrium during the catalytic stage of *S. cerevisiae* pre-mRNA splicing. Following branching, the exon ligation factors Slu7 and Prp18 can bind the C conformation with low affinity by interacting with the Prp8 RNase H domain (Prp8^RH^) to prime remodelling, thus forming the C_i_ intermediate. Prp16-mediated remodelling of this C_i_ complex leads to a C* ‘open’ conformation, with high affinity for Slu7/ Prp18. Dissociation of Cwc25 from C_i_ allows the Yju2 C-terminal domain (Yju2^C^) to interact with Prp8^RH^ and stabilise binding of Prp22. The undocked branch helix is locked in a C* ‘closed’ conformation by the stable binding of Prp18 and Slu7 and by Yju2^C^. The C* closed conformation is competent for stable 3′-SS docking, allowing exon ligation. The C_i_ complex is likely unstable and during Prp16 activity, in the absence of additional Slu7/ Prp18 binding, remodelling can also allow C_i_ to catalyse reversal of branching, thus reverting to a B* conformation in which Slu7/ Prp18 may remain bound (B*^*Slu7/Prp18*^). Remodelling of C complex may also occur through a pathway that does not involve C_i_, where Prp16 action precedes, or is concomitant with, recruitment of Slu7/ Prp18 in the C* ‘open’ conformation.

Consistently, at least 30% of C complexes stalled with the conditional *prp16-302* allele and purified at low salt concentrations contain Prp18 in the C_i_ state (**Extended Data Fig. 8**). These complexes can be reactivated to produce mRNA upon ATP hydrolysis by Prp16 even in the absence of exogenous Slu7 and Prp18 added following purification (**Fig. 4**). We conclude that C_i_ is a key on-pathway intermediate in remodelling between branching and exon ligation (**Fig. 6**): Prp18 and Slu7 pre-bound in C_i_ can stably engage the spliceosome to stabilise the higher-energy C* complex and promote exon ligation following ATP hydrolysis by Prp16 (**Figs. 4d-g**,**6**).

Binding of branching factors is also remodelled during Prp16 activity. In B*/C complexes, branching is promoted by engagement of the N-termini of Cwc25, Yju2, and Isy1 with the branch helix and brA, while the C-termini of these branching factors interact with the NTC and Prp16. Two C-terminal helices of Yju2 cross Cwc25 before interacting with the NTC, forming a unique binding platform for Prp16 at the branching stage^15,53^ (**Fig. 7a-c**). These Yju2 helices are reorganised in C*/P complex to instead bridge the rotated Prp8 RNaseH domain to Prp22, while the Yju2 N-domain dissociates (**Fig. 6d**). While the N-terminus of Yju2 is essential for viability and promotes branching, deletion of the C-terminus allows only inefficient exon ligation in the absence of Prp16 (ref.54), suggesting that Yju2 stabilises both C and C* in a manner consistent with our structural model (**Figs. 6, 7**). Interestingly, we show here that the C-terminus of Isy1 also binds the NTC, gluing the interface between Syf1 and the U2 snRNP (**Fig. 5**). Although detailed structures of the NTC are not yet available in B^act^ and C*/P complexes, this interaction is likely preserved after remodelling to C* complex, while the Isy1 N-domain similarly dissociates. Thus, paralleling the primed association of exon-ligation factors with the branching conformation in C_i_ complex, the branching factors Isy1 and Yju2 are both associated with the exon-ligation conformation in C* and P complex. Indeed, tethering of Isy1 by the NTC throughout catalysis explains how Isy1 affects proofreading of both the branch site and 3′-SS (ref. 13).

**Figure 7.**
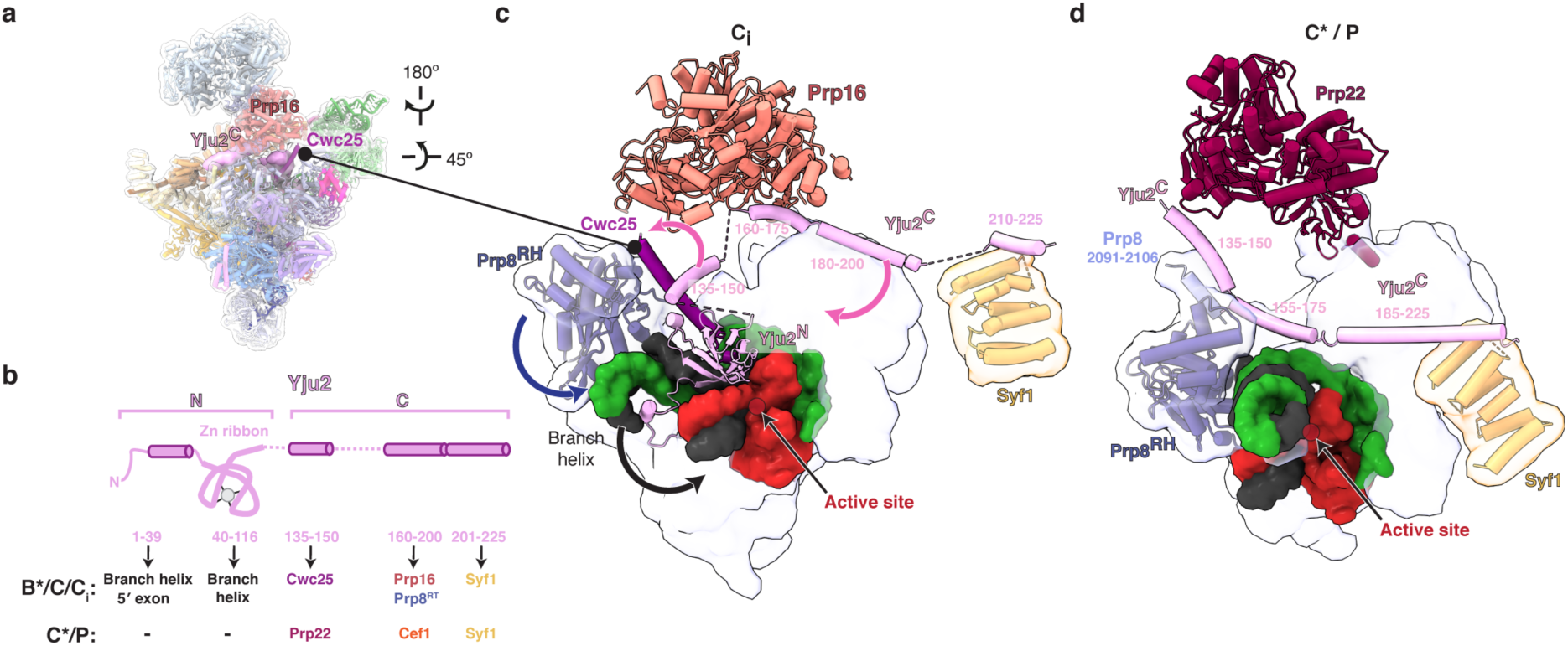
Remodelling of Yju2 interactions during the C to C* transition. **a**, Location of the branching factor Yju2 in the C-complex spliceosome. **b**, Domain architecture of Yju2 and its interaction partners during branching and exon ligation. **c**, Yju2 supports binding of Cwc25 and Prp16 in C complex. **d**, Yju2 is remodelled in C*/P to act as exon-ligation factor. Note that dissociation of Cwc25 is required for formation of a new Prp8 helix that supports Yju2-mediated recruitment of Prp22 in C*/P.

The C_i_ intermediate indicates that complexes in the branching conformation (B*/ C/ C_i_) differ little in composition from those in the exon-ligation conformation (C*/ P). Only Cwc25 dissociates from branching to exon-ligation, while all other factors can already bind in the C conformation and are remodelled as a result of Prp16 action (**Fig. 6**). Indeed, the C conformation is particularly stable, as complexes that cannot complete catalysis appear to revert to this state. C complexes accumulate not only when Prp16 remodelling is blocked but also when exon ligation is impaired by mutation of the 3′-SS, or when mRNA release is prevented by blocking Prp22 activity (**Extended Data Fig. 2**). Therefore, the energy of ATP hydrolysis by Prp16 is necessary to disrupt this stable conformation and establish an equilibrium between branching and exon ligation that underlies reversibility of both splicing reactions^55^ and is likely necessary for proofreading during catalysis.

The C_i_ complex structure shows how exon-ligation factors can prime remodelling before Prp16 activity and shift the equilibrium from branching to exon ligation after Prp16 hydrolyses ATP. By stabilising the higher-energy C* conformation, exon-ligation factors act like a Brownian ratchet, similarly to how branching factors drive the equilibrium towards B* during spliceosome activation by Prp2 (ref. 56, cf. ref. 26). Specific branching factors further modulate this equilibrium through partial dissociation of their N-terminal domains as a result of Prp16 action (**Fig. 6**). Conversely, when Slu7 and Prp18 are limiting, the spliceosome cannot become ‘locked’ into its exon-ligation conformation and Prp16 action therefore allows the C-complex spliceosome to reverse branching and revert to the B* conformation (**Figs. 4d-g, 6**). Such reversibility may be important during proofreading of branching by Prp16 (refs. 26,57) and suggests that Slu7 and Prp18 may affect catalysis and proofreading more broadly, by modulating the equilibrium between B*/ C/ C_i_ and C*/ P complexes. In support of this idea, we observed the C_i_ conformation when spliceosomes were stalled with a 3′-SS mutant that docks poorly in the active site but can produce mRNA when proofreading is disabled^28,58^, suggesting that proofreading of exon ligation by Prp22 can lead to substrate rejection and collapse to the C_i_ conformation without dissociation of exon-ligation factors (**Fig. 6, Extended Data Fig. 8**).

Overall, Prp16 acts as a classical catalyst by reducing the activation barrier to transition from the very stable branching conformation to the higher-energy exon-ligation conformation, thus allowing thermodynamic control of splicing, a feature likely necessary for proofreading of catalysis. Indeed, genetic studies suggest that the stability of RNA elements that form the active site is also disrupted during Prp16 action in a manner that influences splice site proofreading^6,59^.

Our model of the catalytic stage involving the novel C_i_ conformation (**Fig. 6**) explains how the spliceosome uses the energy of ATP hydrolysis to remodel binding of step-specific protein factors and to modulate substrate transactions at a single RNA-based active site, thus allowing conformational equilibrium and proofreading during splicing catalysis.

## Supporting information

PyMol session for the Ci spliceosome

## Data availability

The cryo-EM maps have been deposited in the Electron Microscopy Data Bank with accession codes EMD-NNNNN, EMD-NNNNN, EMD-NNNNN, EMD-NNNNN. The coordinates of the atomic model have been deposited in the Protein Data Bank under accession code XXXX.

For immediate access, the composite map and model, as well as a PyMol session can be downloaded from the following folder, using “Connect As: Guest” if prompted for a password: ftp://ftp.mrc-lmb.cam.ac.uk/pub/mwilkin/Ci_spliceosome/

## Acknowledgements

We thank K. Nguyen for a gift of Slu7/ Prp18 heterodimer used for preliminary tests; S. Scheres, X.C. Bai, C. Savva, S. Chen, K. R. Vinothkumar, and G. Cannone for help and advice on data collection and processing; G. McMullan, J. Grimmett, and T. Darling for smooth running of the EM and computing facilities; the staff at Diamond Light Source (DLS) for help with data collection; A. Amunts and S. Aibara for help with data collection at SciLifeLab; the mass spectrometry facility for help with protein identification; P. Emsley and G. Murshudov for help and advice with model building and refinement; the members of the spliceosome group for help and advice throughout the project. We thank J. Löwe, D. Barford and S. Scheres for their continuing support, C. Charenton, P-C. Lin, S. Lövestam, A. Newman, and S. Routh for critical reading of the manuscript, and J. Staley for a gift of reagents. The project was supported by the Medical Research Council (MC_U105184330) and a European Research Council Advanced grant (SPLICE3D). S.M.F. was supported by EMBO and Marie Skłodowska-Curie fellowships; M.E.W. was supported by a Rutherford Memorial Cambridge Scholarship.

## Author contributions

S.M.F. and W.P.G. established the method for C complex preparation. S.M.F. and M.E.W. established the method for P complex preparation. S.M.F. designed and performed C^*prp16-302*^ experiments including sample and grid preparation and biochemistry. M.E.W. and W.P.G. prepared C complex sample and grids. M.E.W prepared P complex sample and grids. M.E.W., S.M.F., and W.P.G. collected and performed initial processing of EM data. M.E.W. merged and re-processed all datasets, carried out model building and refinement and prepared the data for deposition. M.E.W. and S.M.F. wrote the manuscript with input from all authors. K.N. initiated and coordinated the spliceosome project.

## Methods

### Cloning and protein expression

A TAPS tag cassette was added in frame to the C-terminus of the endogenous CWC25 locus in *S. cerevisiae* yJPS983 (carrying a chromosomal *prp16-302* allele^6^), using the *kanMX6* resistance cassette^60^. *prp16-302* encodes the *prp16R456K* in motif Ib, which significantly reduces ATP hydrolysis at low temperatures and stalls spliceosomes after branching^61^. For extract preparation the resulting strain was grown normally at 30°C in YEPD media in a batch fermenter to an optical density OD_600_ ∼ 2.4 – 3.2.

A construct encoding Prp18 and Slu7, which form a heterodimer^19^, was used for recombinant expression in *E. coli* BL21 (DE3) RIL cells. The expressed heterodimer was purified via Ni-NTA-agarose and gel filtration chromatography.

### C^*prp16-302*^ purification and biochemistry

Spliceosomes were assembled on a modified UBC4 pre-mRNA with 25 nt exons^5^ in extracts from *prp16-302* / *CWC25*-TAPS by incubation at 19°C for 30 minutes. Following splicing, reactions were incubated at 19°C for another 15 minutes in the presence of 0.2 µM of a DNA oligo directed against the 5′-splice site, to degrade unassembled pre-mRNA, and 2 mM glucose to deplete ATP and minimise Prp16 activity. Reactions were centrifuged through a 40% glycerol cushion in buffer K-75 (20 mM HEPES KOH pH 7.9, 75 mM KCl, 0.25 mM EDTA, 0.05% NP-40) and assembled complexes were affinity purified from the cushions using lgG Sepharose 6 Fast Flow (GE Healthcare) to capture the Protein A tag. Following extensive washing with buffer K-75, complexes were eluted from the beads by incubation with 25 ug/ mL TEV protease (expressed in house) by incubation at room temperature (22-23°C) for one hour in the presence of 1 mM DTT, and the protease was removed by subsequent concentration through a 50 kDa MWCO Amicon concentrator. Spliceosomes were further purified via the Strep II tag on Cwc25 using Streptactin affinity resin (GE) in buffer K-75 (20 mM HEPES KOH pH 7.9, 75 mM KCl, 0.25 mM EDTA, 5% glycerol, 0.05% NP-40) and eluted with 5 mM desthiobiotin. Eluted complexes were concentrated in a 100 kDa MWCO Amicon concentrator to ∼ 5-10 nM, buffer exchanged into buffer K-75 without glycerol (20 mM HEPES KOH pH 7.9, 75 mM KCl, 0.25 mM EDTA, 0.0025% NP-40), and used for EM data collection or biochemistry. This sample was used for EM data set 4.

For biochemical assays, purified C complexes were chased at 26°C for 60 minutes in reactions containing 10% concentrated spliceosomes (∼100 fmoles, 2 nM, in K-75 without glycerol), 3% PEG_8000_, 60 mM potassium phosphate buffer (pH 7.0), 2 mM ATP (or ATP-γS), and 4 mM MgCl_2_. Specific reactions were also supplemented with 250 nM Slu7/ Prp18 heterodimer. The data in Fig. 4 were quantified using an Amersham Typhoon imaging system and ImageQuant TL. The rolling ball algorithm was used for background subtraction. Bands for individual splicing species were first normalised to the total signal from all splicing species in each lane to obtain percentages, which were then used for calculating the efficiency of exon ligation and of pre-mRNA formation by reversal of branching from the lariat-intermediate. Quantifications in Fig. 4e,f were obtained from four independent spliceosome purifications using extracts from two independent biological replicates (independent extract preparations from two different batches of yeast fermenter growth).

For EM data collection following biochemical chase, purified C complexes were incubated at 26°C with 2 mM ATP and 2.5 mM MgCl_2_ for 15 minutes prior to grid preparation. This sample was used for EM dataset 5. For both un-chased and chased complexes, the purified sample was applied to Cu 300 R1.2/1.3 holey carbon grids (Quantifoil) coated with a ∼6 nm homemade carbon film. Grids were glow discharged for 30 seconds before application of 3.5 μL sample, then incubated for 25 s and blotted for 2.5-3.5 s before vitrification by plunging into liquid ethane using an FEI Vitrobot MKIII operated at 100% humidity and 4°C.

### P complex purification

Dataset 6 described in this paper derives from our early attempts to purify P complex. mRNA release was stalled by a dominant negative Prp22 mutant (K512A) and spliceosomes were purified first by mRNA pulldown then by Slu7-TAPS pulldown. This strategy produces a mixture of C and C*/P complex and predates our discovery that RNaseH cleavage could be used to significantly enrich P complex in the final sample^9^ (**Extended Data Fig. 1**). In detail, Slu7-TAPS splicing extract was prepared as described^7^. Splicing extracts were treated on ice for 10 min with 0.04 mg/mL recombinantly-expressed Prp22 K512A. Spliceosomes were assembled in this extract on a *UBC4* pre-mRNA with two MS2 hairpins pre-bound to MS2-MBP fusion protein as described^62^ by incubation at 23°C for 30 minutes. Following splicing, reactions were centrifuged through a 40% glycerol cushion in buffer K-75 (20 mM HEPES KOH pH 7.9, 75 mM KCl, 0.25 mM EDTA, 0.05% NP-40) and assembled complexes were affinity purified from the cushions using amylose resin (New England Biolabs). Following washing with buffer K-75, complexes were eluted from the beads with 12 mM maltose. Spliceosomes were further purified via the Strep-II tag on Slu7 using Streptactin affinity resin (GE) in buffer K-100 (20 mM HEPES KOH pH 7.9, 100 mM KCl, 0.25 mM EDTA, 5% glycerol, 0.01% NP-40) and eluted with 5 mM desthiobiotin. Eluted complexes were concentrated in a 100 kDa MWCO Amicon concentrator and dialysed into buffer K-75 without glycerol (20 mM HEPES KOH pH 7.9, 75 mM KCl, 0.25 mM EDTA). This sample was used for EM dataset 6: the purified sample was applied to Cu 300 R1.2/1.3 holey carbon grids (Quantifoil) coated with a ∼7 nm homemade carbon film. Grids were glow discharged for 30 seconds before application of 3 μL sample, then incubated for 30 s and blotted for 2.5 s before vitrification by plunging into liquid ethane using an FEI Vitrobot MKIII operated at 100% humidity and 4°C.

### Cryo-EM data acquisition

Eight data sets and a total of 24,115 movies were collected manually or using EPU on various Titan Krios microscopes (Thermo Fisher) all equipped with energy filters (slit width of 20 eV) and K2 detectors operated in counting or super-resolution mode (**Extended Data Fig. 2**). Dataset 1 (2,213 movies) is described^7^ and was collected manually on LMB Krios 1 in super-resolution mode with a real pixel size of 1.427 Å/pix, a defocus range of −0.5 to −4 μm, with an exposure time of 16 s fractionated into 20 frames and a total dose per micrograph of 40 e^−^/Å^2^. Datasets 2 and 3 were described previously^8^. Dataset 2 (1,571 movies) was collected manually on LMB Krios 1 in super-resolution mode with a real pixel size of 1.427 Å/pix, a defocus range of −0.5 to −4.5 μm, with an exposure time of 16 s fractionated into 20 frames and a total dose per micrograph of 40 e^−^/Å^2^.

Dataset 3 (2,025 movies) was collected using EPU on Diamond Light Source (DLS) Krios 1 in counting mode with a pixel size of 1.031 Å/pix, a defocus range of −0.5 to −3.5 μm, with an exposure time of 14 s fractionated into 28 frames and a total dose per micrograph of 42 e^−^/Å^2^.

Dataset 4 (2,944 movies) was collected using EPU on Diamond Light Source (DLS) Krios 2 in counting mode with a pixel size of 1.023 Å/pix, a defocus range of −0.5 to −3.5 μm, with an exposure time of 14 s fractionated into 28 frames and a total dose per micrograph of 44 e^−^/Å^2^.

Dataset 5 (2,523 movies) was collected using EPU on LMB Krios 1 in counting mode with a pixel size of 1.12 Å/pix, a defocus range of −0.5 to −3 μm, with an exposure time of 14 s fractionated into 28 frames and a total dose per micrograph of 56 e^−^/Å^2^

Dataset 6 (4,369 movies) was collected using EPU on a Krios at SciLifeLab (Stockholm, Sweden) in counting mode with a pixel size of 1.028 Å/pix, a defocus range of −0.5 to −3.5 μm, with an exposure time of 8 s fractionated into 20 frames and a total dose per micrograph of 39.7 e^−^/Å^2^.

Dataset 7 (2,384 movies) is described in ref. 9 and was collected using EPU on LMB Krios 1 in counting mode with a pixel size of 1.12 Å/pix, a defocus range of −0.2 to −3 μm, with an exposure time of 12 s fractionated into 20 frames and a total dose per micrograph of 47 e^−^/Å^2^.

Dataset 8 (1,614 movies) is described in ref. 63 and was collected using EPU on LMB Krios 1 in counting mode with a pixel size of 0.88 Å/pix, a defocus range of −0.5 to −3.5 μm, with an exposure time of 7 s fractionated into 35 frames and a total dose per micrograph of 45.2 e^−^/Å^2^.

Dataset 9 was of a P complex sample prepared identically to ref. 9 except that the final sample buffer contained 1 mM MgCl_2_ instead of 0.25 mM EDTA. 4830 micrographs were collected using EPU on a Diamond Light Source Krios in counting mode with a pixel size of 1.03 Å/pix, a defocus range of −0.5 to −3 μm, with an exposure time of 12 s fractionated into 40 frames and a total dose per micrograph of 49.2 e^−^/Å^2^.

### Initial cryo-EM data processing

All datasets were initially processed separately (**Extended Data Fig. 2**). Dataset 1 was processed as described previously^7^ yielding a 3.8 Å reconstruction of C complex (EMD-4055). Re-refining these particles with a mask on the spliceosome core improved the resolution to 3.61 Å. Datasets 2 and 3 were processed as described previously^8^, yielding both a reconstruction of C* complex at 3.8 Å resolution (EMDB-3539) and reconstructions of C complex at 4.62 Å resolution (dataset 2) and 3.61 Å resolution (dataset 3).

Datasets 4 and 5 were processed similarly: particles were picked using RELION auto-pick using representative 2D class averages of C complex as templates. After 3D classification to select good quality particles, particles were polished using the method implemented in RELION 1.3 (ref. 64, i.e. not the Bayesian polishing routine of RELION 3.0). Masked refinement then yielded C complex reconstructions at 3.34 Å resolution (dataset 4) and 3.37 Å resolution (dataset 5).

Dataset 6 was processed similarly to datasets 4 and 5 except that due to splicing being stalled prior to mRNA release, approximately half the particles were in the exon-ligation conformation (C*/P) and half were in the branching conformation (C). 3D classification was used to resolve these different populations, and each was subjected to particle polishing in RELION 2.0, producing a 3.58 Å resolution reconstruction of C*/P complexes, and 3.42 Å resolution reconstruction of C complex.

Dataset 7 (ref. 9) was reprocessed in RELION 3.1. After motion-correction with dose-weighting, particles were picked with crYOLO (ref. 65) using a model trained on the dataset. After 3D classification to select good P complex particles, particles were subject to CTF refinement and Bayesian polishing, producing a P complex reconstruction at 3.50 Å resolution.

Dataset 8 was processed as described in ref. 63, yielding a 3.56 Å reconstruction of P complex.

Dataset 9 processed in RELION 3.1. After motion-correction with dose-weighting, particles were picked with crYOLO (ref. 65) using a general model. After 3D classification to select good P complex particles, particles were subject to CTF refinement and Bayesian polishing, producing a P complex reconstruction at 3.13 Å resolution.

### C complex data processing

#### Initial refinement

C complex data were merged using RELION 3.1, using separate optics groups for datasets 1, 2, 3, 4, 5, and 6 (**Extended Data Fig. 2**). Dataset 1 was further split into 3 optics groups as this dataset was collected over three different microscope sessions. The individual reconstructions were compared with UCSF Chimera to determine the relative pixel sizes of each dataset, and defocus values were scaled by the relative difference in pixel sizes^63^. Unscaled particles were then refined together with a mask on the ordered core, with RELION 3.1 internally scaling the particles to match a 1.12 Å/pix reference in a 400 pixel box. Two rounds of CTF refinement (refining per-particle defocus, per-micrograph astigmatism and B-factor, and per-optics group anisotropic magnification, beam tilt, trefoil, and 4^th^ order aberrations) produced a reconstruction at 2.80 Å resolution (**Extended Data Fig. 4**). The reconstruction had a good angular distribution that was improved by merging the 6 datasets (**Extended Data Fig. 4**). Density modification in Phenix (ref. 66) further improved the map resolution to 2.69 Å.

#### Focussed refinements

Although the core of the C-complex spliceosome is well ordered, many of the peripheral domains are flexible and had weak EM density in the overall refinement map, and correspondingly the local resolution quickly decayed towards the edge of the spliceosome (**Extended Data Fig. 4**). For each domain, strategies including signal subtraction, classification without alignment, and focussed refinement with and without the reconstruction algorithm SIDESPLITTER (ref. 67) were systematically investigated. All signal subtractions started from a 3 Å resolution map of C complex where the refinement mask encompassed the entire complex. All focussed refinements used solvent-flattened FSCs to calculate the resolution during refinement. The following strategies produced the best maps for each domain, and are summarised in **Extended Data Fig. 3** and **Table 1**.

Density for the U2 snRNP core domain was improved first by signal subtraction with a soft mask (8 pixel hard edge, 16 pixel soft edge) encompassing the U2 snRNP core, Prp8 RNaseH domain and Syf1 N-terminus. Signal-subtracted particles were shifted to the mask centre-of-mass and cropped to a 200 pixel box. 3D classification without alignment and T=4 into 4 classes required 60 iterations for convergence and showed that 26% of particles (108,540) did not have strong U2 snRNP density. These particles were removed, and the resultant STAR file was reverted to the original, un-subtracted particles. These were focus-refined using the same mask used for signal subtraction, using the overall map low-pass filtered to 7 Å as a reference and performing local angular searches starting at 0.9 degree sampling. This produced a reconstruction at 4.07 Å resolution. Refinement was then continued using a tighter mask that only encompassed the U2 snRNP core (3 pixel hard edge, 16 pixel soft edge), using a 5 Å resolution reference and performing local angular searches at 0.5 degree sampling. This produced a map for the U2 snRNP at 3.58 Å resolution. Density modification in Phenix (ref. 66) further improved the map resolution to 3.26 Å.

The NTC TPR domains (Syf1 and Clf1 and associated proteins) was improved first by signal subtraction with a mask (1 pixel hard edge, 8 pixel soft edge) encompassing the entirety of Clf1, Syf1 and the associated proteins. Signal-subtracted particles were shifted to the mask centre-of-mass and cropped to a 300 pixel box. 3D classification without alignment and T=4 into 4 classes required 25 iterations for convergence and showed that 43% of particles (172,895) did not have strong density for the peripheral helical arches. These particles were removed, and the remaining particles were focus-refined using a soft mask (8 pixel hard edge, 16 pixel soft edge) that only encompassed the flexible peripheral helical arches, e.g. excluding the well resolved N-terminus of Clf1. The best 3D class, low-pass filtered to 8 Å, was used as a reference, local angular searches started at 0.9 degree sampling, and the external reconstruction program SIDESPLITTER was used to reduce overfitting. This produced a reconstruction at 3.82 Å resolution. Refinement was then continued for 3 more iterations with the same mask, using a 3.8 Å resolution reference and performing local angular searches with 0.9 degree sampling. This produced an improved reconstruction with smoother densities, although the resolution stayed constant at 3.82 Å. Density modification in Phenix (ref. 66) then further improved the map resolution to 3.50 Å.

Despite numerous attempts, the helicase module of C complex, consisting of Brr2, Prp16, and the Prp8 Jab1/MPN domain, could not be improved to near-atomic resolution. This module is larger than the U2 snRNP core which was successfully refined to near-atomic resolution. Therefore, this module may have internal flexibility, e.g. between the RecA domains of Brr2 and Prp16, that makes particles difficult to align. Nevertheless, densities were made more interpretable as follows. First signal subtraction was performed using a mask (1 pixel hard edge, 8 pixel soft edge) loosely encompassing Brr2, Prp16, and Prp8 Jab1/MPN. Signal-subtracted particles were shifted to the mask centre-of-mass and cropped to a 300 pixel box. 3D classification without alignment and T=4 into 4 classes required 60 iterations for convergence and showed that only 26% of particles (103,461) had strong density for the helicase module. These were focus refined with a very soft mask (16 pixel hard edge, 12 pixel soft edge), using 0.9 degree local angular sampling, and the best class low-pass filtered to 20 Å as a reference. Postprocessing with a tighter mask (1 pixel hard edge, 16 pixel soft edge) gave a reconstruction of the helicase module at 8 Å resolution, which was used for interpreting Prp16. Classification without alignment suggested that Brr2 and Prp8 Jab1/MPN was flexible relative to Prp16. Therefore, a further focussed refinement was performed using a soft mask (8 pixel hard edge, 16 pixel soft edge) just around Brr2 and Prp8 Jab1/ MPN, with a 20 Å resolution reference, 0.9 degree local angular sampling, and an initial offset search range of 3 pixels, producing a map at 7.15 Å resolution. Similar attempts to perform focussed refinement on just Prp16 were not successful.

In the overall C-complex map, the WD40 domain of Prp17 was weakly defined and did not allow unambiguous docking of the model from P complex, with all 14 possible orientations of the 7-bladed beta propeller being consistent with the density. However, this domain was too small for focussed refinement. Instead, to improve the local resolution we performed multiple rounds of classification without alignment and with high T values to select for high-resolution subsets of particles with Prp17 in similar positions. First, signal subtraction was performed using a loose spherical mask (12 pixel hard edge, 8 pixel soft edge) around the WD40 domain and some neighbouring proteins. Signal-subtracted particles were shifted to the mask centre-of-mass and cropped to a 200 pixel box. 3D classification without alignment and T=50 into 4 classes required 60 iterations for convergence. The single class with the most well-defined density (24%, 96,800 particles) was selected and 3D classification without alignment was repeated using a tighter mask (1 pixel hard edge, 8 pixel soft edge), T=50, and 4 classes. Again, the class with best defined density (58%, 59,152 particles) was selected. The selected particles were then reverted to the un-subtracted particles, and half maps were reconstructed from the original Euler angles using relion_reconstruct. Postprocessing with a mask encompassing the entire C complex produced a map at overall 3.67 Å resolution. The local resolution around the Prp17 WD40 domain was still limited, but now allowed unambiguous docking of the model from P complex, with one of the 14 possible orientations giving clearly higher correlations and atom inclusion scores when docking in UCSF Chimera.

#### Focussed classification of step II factors

To obtain robust estimates of the occupancy of the step II factors Prp18 and Slu7 on the C complex spliceosome, we used multiple parallel 3D classifications and merged selected classes (**Extended Data Fig. 3**). To identify particles containing the Slu7 zinc knuckle domain, signal subtraction was used with a small spherical mask (8 pixel hard edge, 8 pixel soft edge) centred on Slu7. Signal-subtracted particles were shifted to the mask centre-of-mass and cropped to a 200 pixel box. Two parallel 3D classifications without alignment were performed, both into 4 classes with 60 iterations and using a tighter spherical mask (1 pixel hard edge, 8 pixel soft edge). One classification used T=100, the other used T=1000. The T=100 classification showed 29% of particles contained the zinc knuckle domain, and the T=1000 classification showed that 18% of particles contained the same domain. After merging these particles and removing duplicates, 33% (130,588) of the original particles were captured, and further classification without alignment did not result in further segregation into different classes: using T values ranging from 100 to 2000, 85 – 95% of particles would converge on a single class with strong zinc knuckle density. Therefore, these 33% were taken as the final “Slu7 zinc knuckle” class.

To identify particles containing Prp18 and the associated EIE region of Slu7, 3D classification was performed both with varying T values and with and without signal subtraction. Signal-subtracted particles were generated using a soft mask centred on Prp18 (4 pixel hard edge, 12 pixel soft edge), and were shifted to the mask centre-of-mass and cropped to a 200 pixel box. Two parallel 3D classifications without alignment were performed on the signal-subtracted particles, both into 4 classes and using the same mask as for signal subtraction. One classification used T=4 and 60 iterations, the other used T=10 and 120 iterations. The T=4 classification showed 20% (82,492) of particles with strong density for Prp18, and the T=10 classification showed that 20% (81,431) of particles contained the same domain. These subsets were merged and duplicate particles were removed, yielding 23% (91,424) of the total particles.

Two parallel 3D classifications without alignment were also performed on the non-signal-subtracted particles, both into 4 classes with 60 iterations and using the same mask as for the signal-subtracted particles. One classification used T=4 and showed 20% (82,398) of particles with strong Prp18 density, the other classification used T=20 and showed 21% (82,872) of particles with strong Prp18 density and 21% (83,389) with some weak Prp18 density. These subsets were merged with the 91,424 un-subtracted particles from the signal-subtracted classifications, and duplicate particles were merged, yielding 186,323 particles. These were subjected to a final 3D classification without alignment into 4 classes, using a looser mask (8 pixel hard edge, 8 pixel soft edge), 60 iterations, and T=20. Two of the resulting classes had strong Prp18 density and were selected as the final “strong Prp18” class, containing 120,141 particles, or 30% of the original particles.

Contingency tables for each dataset (**Extended Data Fig. 8**) were determined from the above classifications, i.e. the datasets were not individually classified.

### Merging C and P complex data to improve Prp19, Cwc22, and U5 Sm reconstructions

The Prp19 module, U5 Sm domain, and the N-terminal domain of Cwc22, are each very flexible but are in similar positions in C and P complexes as they are unaffected by the conformational change between step I and step II of splicing. We therefore reasoned that the best resolutions for these regions could be obtained by merging C and P complex data to give high initial particle numbers for focussed classification.

First, all P complex data were first merged using RELION 3.1, using separate optics groups for datasets 6, 7, 8, and 9 (**Extended Data Fig. 2**). The individual reconstructions were compared with UCSF Chimera to determine the relative pixel sizes of each dataset, and defocus values were scaled by the relative difference in pixel sizes^63^. Unscaled particles were then refined together, with RELION 3.1 internally scaling the particles to match a 1.12 Å/pix reference in a 400 pixel box. Two rounds of CTF refinement (refining per-particle defocus, per-micrograph astigmatism, and per-optics group anisotropic magnification, beam tilt, trefoil, and 4^th^ order aberrations) followed by a final masked refinement produced a reconstruction at 2.95 Å resolution from 320,770 particles. This map had unclear density for the docked 3′ splice site, so we still consider our previous reconstruction at 3.30 Å with strong 3′ splice site density as the best “reference” P complex structure^63^.

Next, we merged all our C complex particles with these P complex particles described above, giving a total of 724,244 particles that when refined produced a map that largely resembled C complex (presumably due to the majority of the particles – 56% – being in the step I conformation) although with weaker density for the mobile U2 snRNP, the branch helix, and the step I factors (**Extended Data Fig. 3**). For the extremely mobile Prp19 domain of the NTC, no focussed refinements were successful. We therefore performed signal subtraction with a loose mask that would encompass most possible orientations of this domain, constructed by summing all classes from a preliminary classification without alignment, then adding a 2 pixel hard edge and 8 pixel soft edge. Signal-subtracted particles were shifted to the mask centre-of-mass and cropped to a 300 pixel box. 3D classification without alignment and T=70 into 12 classes showed there were indeed many possible orientations of this domain (**Extended Data Fig. 3**). The most stable class, containing 7% of the original particles (49,514) was selected. Half maps were reconstructed for the signal-subtracted particles using relion_reconstruct and the original Euler angles, allowing postprocessing with a soft mask (hard edge 8 pixels, soft edge 16 pixels) encompassing just the stable orientation, which produced a map at nominally 7.30 Å resolution. This resolution is probably overestimated but the map nonetheless shows some secondary structure features consistent with the expected helical bundle (**Extended Data Fig. 5**). We also reverted to the original C-complex particles (35,716) and calculated half-maps with relion_reconstruct to show how this stable orientation relates to the body of the spliceosome. After postprocessing with a mask around the entire C-complex, this produced a reconstruction at 3.97 Å resolution (local resolution around Prp19 ∼ 8 Å) showing how the N-terminus of Prp46 probably projects into the Prp19 module.

A similar approach was applied for the U5 Sm domain (**Extended Data Fig. 3**), as this domain was too small for focussed refinements. Signal subtraction was performed on the combined C/C*/P particles using a soft mask encompassing the U5 Sm ring and the base of U5 snRNA (hard edge 4 pixels, soft edge 8 pixels), and re-centering particles on the mask centre-of-mass and cropping to a 200 pixel box. Two rounds of 3D classification without alignment were used to select the particles with the most stably-bound, highest resolution U5 Sm domain. The first round with T=20 eliminated 54% of particles which showed no density for the U5 Sm domain, and the second round with T=100 selected 23% of particles (65,561 particles, 9% of original particles) which were used to calculate half-maps with relion_reconstruct using the original Euler angles. After postprocessing with a mask around the entire spliceosome, this produced a reconstruction at 3.13 Å resolution with much stronger density for the U5 Sm ring than the original map. The map was further improved by density modification in Phenix (ref. 66) to 2.96 Å resolution, which smoothened some of the discontinuous densities in the peripheral Sm chains.

Finally, a similar approach was utilised for the N-terminal domain of Cwc22 (**Extended Data Fig. 3**). Signal subtraction was performed on the combined C/C*/P particles using a soft mask encompassing the Cwc22 NTD (hard edge 4 pixels, soft edge 8 pixels), and re-centering particles on the mask centre-of-mass and cropping to a 200 pixel box. Two rounds of 3D classification without alignment were used to select the most stable, high-resolution position of Cwc22 NTD. The first round with T=100 eliminated 72% of particles which showed no density for Cwc22 NTD, and the second round with T=1000 selected 46% of particles (73,802 particles, 10% of original particles) which were used to calculate half-maps with relion_reconstruct using the original Euler angles. After postprocessing with a mask around the entire spliceosome, this produced a reconstruction at 3.16 Å resolution. The map was further improved by density modification in Phenix (ref. 66) to 2.97 Å resolution.

### C complex model building

To avoid being biased by our previous model of C complex^7^ which was built into lower resolution density (3.8 Å average resolution), the entire ordered core of C complex was rebuilt entirely de novo in Coot (ref.68), based only on the 2.8 Å cryoEM density. This included the proteins Prp8 (except the RNaseH and Jab1/MPN domains), Snu114, Prp45, Prp46, Ecm2, Cwc2, Cwc15, Bud31, Cef1, Clf1, Syf1, Syf2, Cwc21, Cwc22, Yju2, Cwc25, Isy1, and the U2, U5, and U6 snRNAs, the intron, and the 5′ exon. To improve model geometry, we compared the scale of the map to crystal structures of Prp8 (PDB 4I43, ref.69) and the NMR structure of Bud31 (PDB 2MY1, ref.70), which showed that the true pixel size of the map should be 1.145 Å/ pix. The model coordinates were all scaled by 1.145/1.12 = 1.02232 before further model building.

ISOLDE (ref.71) was then used to diagnose and fix errors and improve Ramachandran, CaBLAM, and rotamer outliers. The resultant core model largely resembled our earlier model, but with more accurate backbone geometry and rotamer assignment (**Table 1**), and some fixing of register errors including in Cef1 helix residues 230-249 and the Cwc15 N-terminus. Notable areas of improvement include the C-terminal domain of Ecm2, the interface of U2 snRNA stem IIb with Cwc2, domain IV of Snu114, assignment of the C-terminal helices of Yju2, and assignment of the very N-terminus of Prp45 and an N-terminal helix of Prp46 projecting towards Prp19.

For building into the focussed-refined maps, these maps were first all aligned and resampled in Chimera to their average positions, as determined from their most populated 3D classes. phenix.combine_focused_maps was then used to make a composite map from the resampled, density-modified maps for the core (2.69 Å), U2 snRNP (3.26 Å), NTC (3.50 Å), U5 Sm (2.96 Å) and Cwc22 NTD (2.97 Å), and models were built into this composite map.

The helical arches of Clf1 and Syf1 were built de novo into the NTC part of the composite map using Coot. The map was of sufficient quality to allow unambiguous building of the N-terminal half of Clf1 and the middle region of Syf1 (**Extended Data Fig. 5**). However, the local resolution still deteriorated towards the periphery of the focus-refined map (**Extended Data Fig. 4**), so we used secondary structure predictions generated by the GeneSilico MetaServer (ref.72) and cross-linking data from yeast B^act^ complex (ref. 73) to support our building. Multiple sequence alignment was also useful, revealing a yeast-specific insertion in Syf1 that we predicted formed a disordered loop, since the cryoEM density of human Syf1 in human P complex (ref. 74) largely resembled the cryoEM density for yeast Syf1. The completed Syf1 model left several helical densities unassigned, with no topological means of filling them with Syf1 sequence. Based on further secondary structure predictions, crosslinking in yeast B^act^ complex, and in some cases clear side-chain density, we assigned these densities to Ntc20, Isy1, and Syf2. This left two remaining helices. One was tentatively assigned to Cef1 but was built with UNK residues, and the other is left unassigned (chain X, UNK residues). The model was then improved in (ref. 71).

The Sm domain of the U2 snRNP was built by docking in the structure of the yeast U1 snRNP Sm domain from pre-B complex (ref. 75) and manually adjusting all side chains and loops to fit the density using Coot. Homology models for Lea1 and Msl1 were generated with I-TASSER (ref.76) based on the structure of human U2A′/U2B′′ in complex with U2 snRNA (PDB 1A9N, ref.77) and were docked into the density and manually fixed. Density connecting the U2 snRNP to Syf1 could not be accounted for by Syf1 helices, and was assigned to Isy1 and the C-terminus of Lea1 based on side-chain density and crosslinking data from yeast B^act^ complex. Double-stranded RNA density corresponding to stems IV, V and the 950 nt yeast-specific insertion in U2 snRNA, was modelled by generating idealised structures in RNAComposer (ref. 78) and adjusting to the density in Coot. The model was then improved using ISOLDE, with adaptive distance restraints used to maintain pairing in U2 stems IV and V.

The Sm domain of U5 snRNP was built by copying our new model for the U2 snRNP Sm domain and docking into the composite map. The associated region of U5 snRNA was docked from yeast B complex (ref.79). The fit for both was improved used using ISOLDE, with adaptive distance restraints on the Sm ring using U2 snRNP as a template, and strong adaptive distance restraints (kappa value of 50) on U5 snRNA.

The Cwc22 N-terminal domain was built by docking an I-TASSER model based on the crystal structure of human Cwc22 complexed with eIF4AIII (PDB 4C9B) into the composite map. Extraneous loops were truncated and the fit was improved with ISOLDE.

Prp17, the Prp8 RNaseH domain, Prp18, and Slu7 all have stronger densities in P complex than in C complex. These were therefore rebuilt using ISOLDE into our newer 3.3 Å map (EMD-10140, ref.63), scaled up to 1.145 Å/pix from 1.12 Å/pix, starting from our original deposited 3.7 Å P-complex model (PDB 6EXN, ref. 9). Prp17, Slu7 zinc-knuckle, and Prp8 RNaseH + Prp18 + Slu7 EIE region were each docked into their respective focus-classified C-complex maps, and ISOLDE was used to fix any clashes and improve the fit to density.

For the helicase domain, a composite map was prepared with phenix.combine_focused_maps from the overall helicase focussed refinement, which had best density for Prp16, and the Brr2 focussed refinement, which had better density for Brr2 and Prp8 Jab1. The crystal structure of Brr2 associated with the Jab1/MPN domain of Prp8 (PDB 4BGD, ref.80) was docked into this composite map after minor rebuilding into the original structure factors with ISOLDE and refinement with phenix.refine to improve starting geometry. A homology model of Prp16 was generated with I-TASSER, with the top two templates being Prp43 (PDB 3KX2) (ref. 81) and MLE helicase (PDB 5AOR), and was docked into the composite map. ISOLDE with adaptive distance restraints was used to morph these starting models into the density. Very strong restraints (kappa of 20) were used for Brr2/Jab1 since the starting model was known to be of good quality. For Prp16, the individual domains (RecA1, RecA2, and CTD) were individually restrained with kappa of 10 to allow movement between them. The resulting Brr2 model was largely unchanged, but Prp16 adopted an open conformation that strongly resembled the crystal structure of *Chaetomium thermophilum* Prp22 in complex with RNA (PDB 6I3P, ref. 82). Indeed, equivalent density to the Prp22-bound RNA was observed next to the RecA2 domain, which based on distance to the modelled intron in the spliceosome core was assigned as residues 87 – 92 of the UBC4 intron (+17 to +22 relative to the branch point, −9 to −4 relative to the 3′ splice site). Prp16 and this region of the intron were then further refined in ISOLDE using strong adaptive distance restraints to the *Ct*Prp22-RNA crystal structure. Finally, the density did not allow correction of Ramachandran and CaBLAM outliers which were inherited from the source structures, but all rotamer outliers were manually corrected when possible.

The Prp19 module, consisting of the Prp19 tetramer, Snt309, and Cef1 C-terminus, was modelled by first rebuilding our earlier model of the human orthologue in human P complex (ref.74) using ISOLDE. An extra helix was built as chain X, residues UNK, but is likely the N-terminus of Prp46. Multiple sequence alignments were then used to mutate and truncate this model to fit the yeast sequences using phenix.sculptor. The U-box domains of Prp19 were modelled using crystal structures of the yeast protein (PDB 2BAY, ref. 83). This model was then docked into the 7.3 Å focussed classification maps of the Prp19 module. This map was good enough to allow improvement of the fit using ISOLDE, using adaptive distance restraints to the starting positions, except for the U-box domains which were strongly restrained (kappa=50) to the crystal structure positions. This fitting showed a pronounced bend in one of the Prp19 dimeric coiled-coils compared to the human structure. After fitting, a large portion of unassigned density remained between the Prp19 module and Syf1. Although low resolution, this density neatly accommodates the crystal structure of the Prp19 WD40 domain (PDB 3LRV, ref.83), which was docked in after rebuilding in ISOLDE and refinement into the original structure factors using phenix.refine.. The Prp19 tetramer should have four such domains, and previous cryoEM structures have shown these in a wide variety of positions, generally without strong support from density. The current assignment of the WD40 domain should also be viewed as tentative, although more well-supported than previous assignments.

### C complex model refinement

Coordinates for the core of C complex, U2 snRNP, NTC, U5 Sm, and Cwc 22 N were refineds imultaneously in real space using phenix.real_space_refine in PHENIX (ref. 84) into a composite map of the 5 respective density-modified focussed maps. Base-pairing, base-stacking, and metal-coordination restraints were not imposed for the core, where the density was good enough to obtain better geometry and fit to density in the absence of external restraints. For refinement of the lariat-intron 2’-5’ linkage a custom set of restraints was adapted from the geometry of the 3’-5’ phosphodiester RNA backbone. Base-pairing and base-stacking restraints were however imposed for U5 snRNA near the U5 Sm site and U2 snRNA stems IIb/c, IV and V. We found that the default real_space_refine settings consistently worsened the excellent starting geometry of our models from ISOLDE. Systematic investigation found that performing one macro-cycle of global minimisation and ADP refinement, skipping local grid searches, with a nonbonded weight of 2000 and overall weight of 0.5, gave the best model quality statistics (**Table 1**). The Prp19 module, helicase module, and Prp17 WD40 domain were not refined in PHENIX, but ISOLDE was used to resolve any clashes between these modules and the refined structure.. The final C complex model comprises 45 protein chains, 3 snRNAs, and the intron and 5′ exon.

### Model visualisation

All structural figures were generated with UCSF ChimeraX (ref. 85).

---

**Extended Data Fig. 1.**
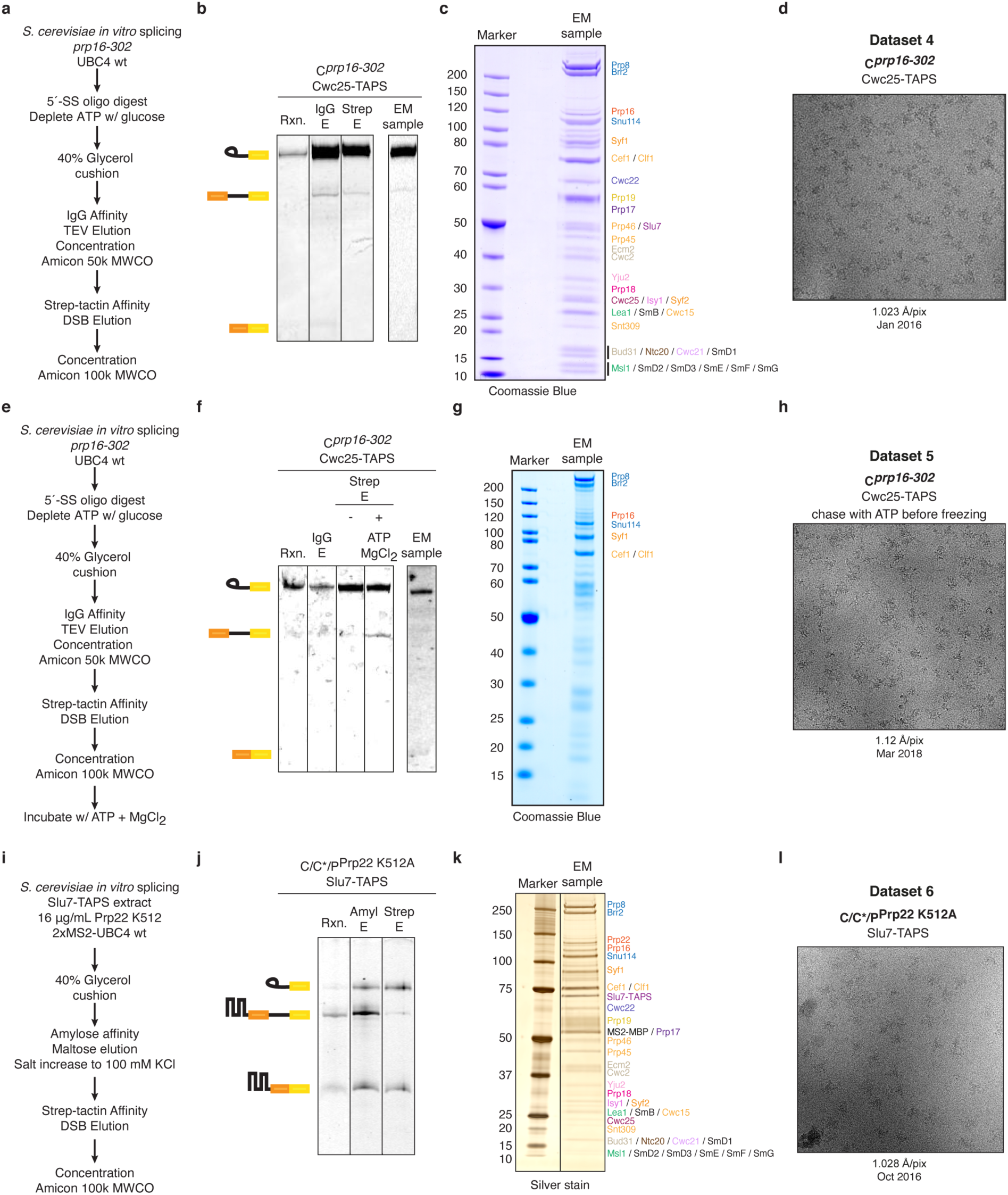
Purification and cryo-EM of new spliceosome samples used for this study. **a-d**, Purification scheme, RNA and protein composition, and representative micrograph for C-complex Dataset 4. **e-h**, Purification scheme, RNA and protein composition, and representative micrograph for C-complex Dataset 5. **i-l**, Purification scheme, RNA and protein composition, and representative micrograph for the P-complex Dataset 6.

**Extended Data Fig. 2.**
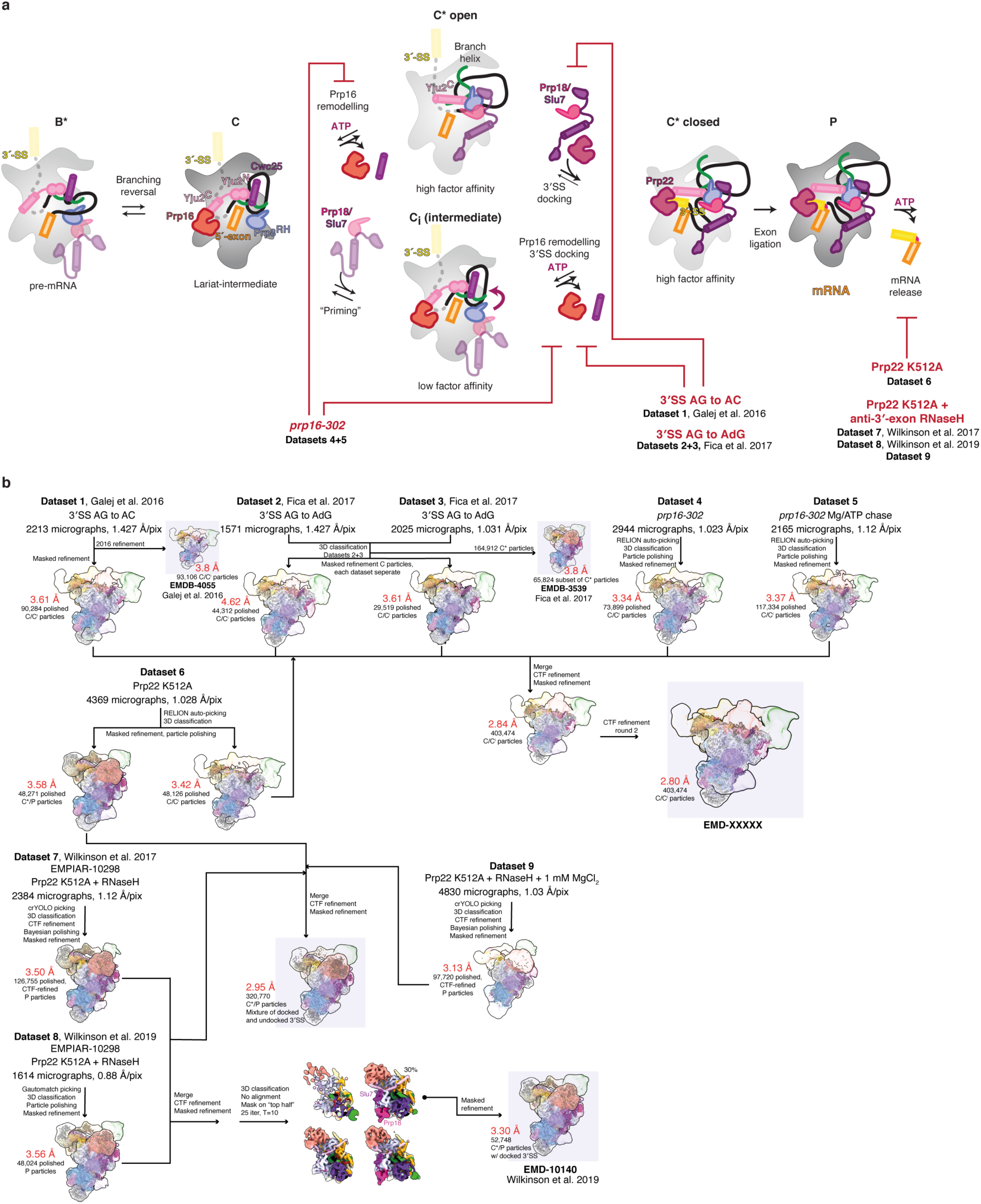
Cryo-EM data processing scheme for dataset merging. **a**, Biochemical scheme for stalling of spliceosomes used for specific datasets. **b**, Dataset merging scheme for producing overall C- and P-complex structures. Final maps deposited to the EMDB are highlighted.

**Extended Data Fig. 3.**
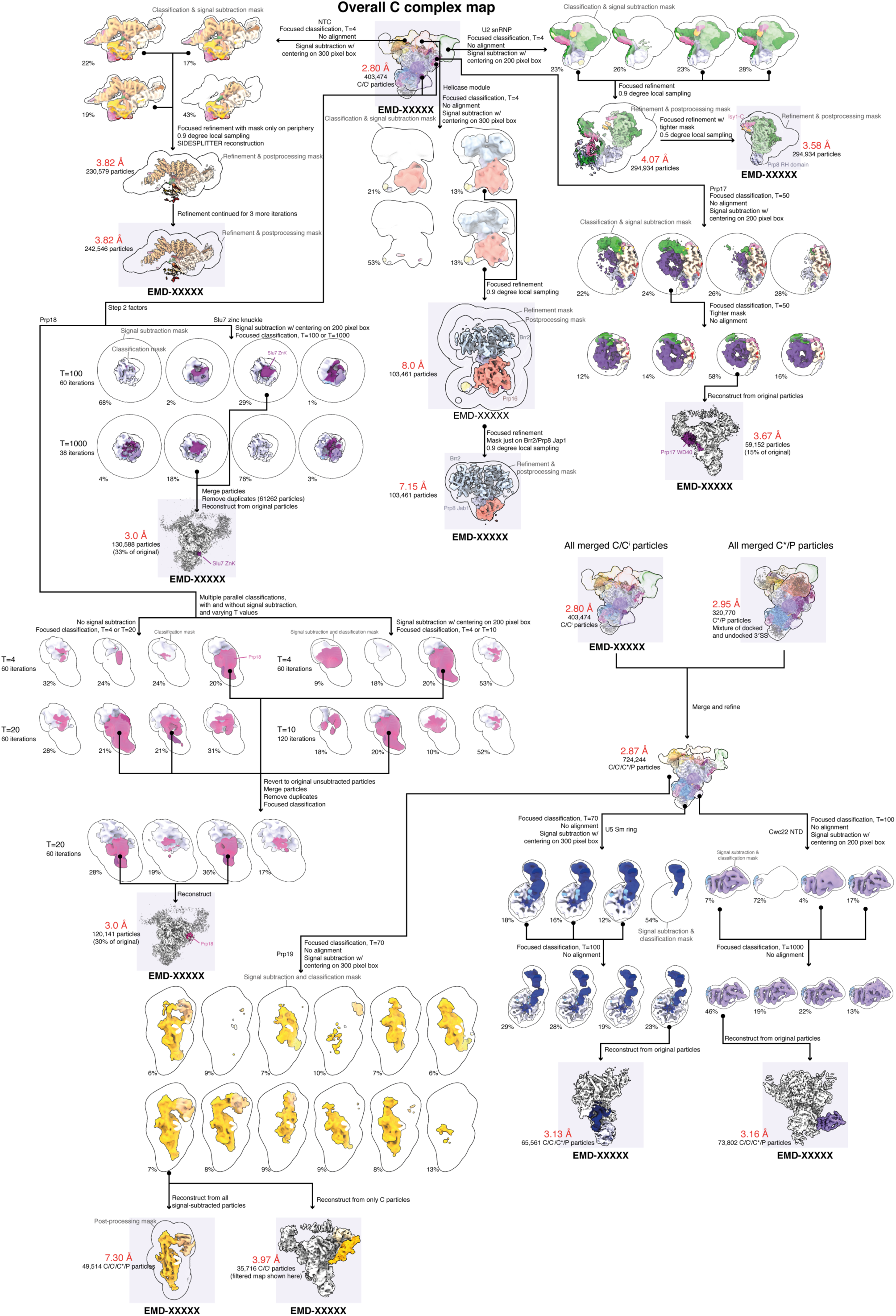
Focussed classification scheme for peripheral regions of the C complex. Final maps deposited to the EMDB are highlighted.

**Extended Data Fig. 4.**
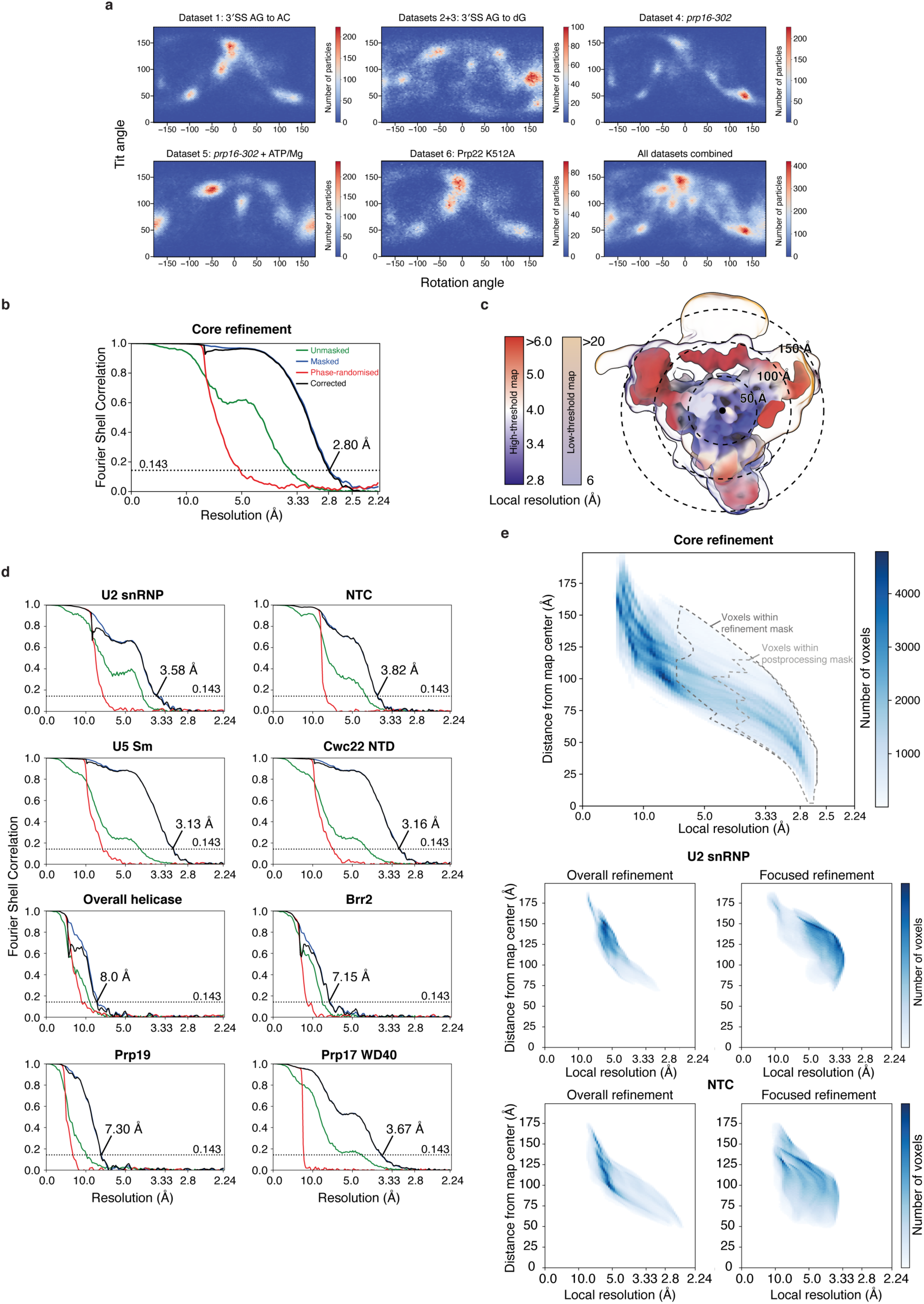
Statistics for the cryo-EM datasets and the resulting maps. **a**, Orientational distribution for the C-complex cryo-EM datasets. Merging datasets improves the overall angular distribution. **b**, Gold-standard Fourier-Shell Correlation curves for the overall C-complex reconstruction and the focus-refined or classified maps. **c**, Overall C-complex reconstruction coloured by local resolution. The map is shown at two thresholds coloured with different palettes to show the high core resolution and low peripheral resolution. **d**, Heatmap of local resolution against distance from map center, showing how in the overall C-complex reconstruction the local resolution decreases towards the map periphery. **e**, Local resolution heatmaps for the U2 snRNP and NTC before and after focussed refinement. Focussed refinement improves the resolution distribution.

**Extended Data Fig. 5.**
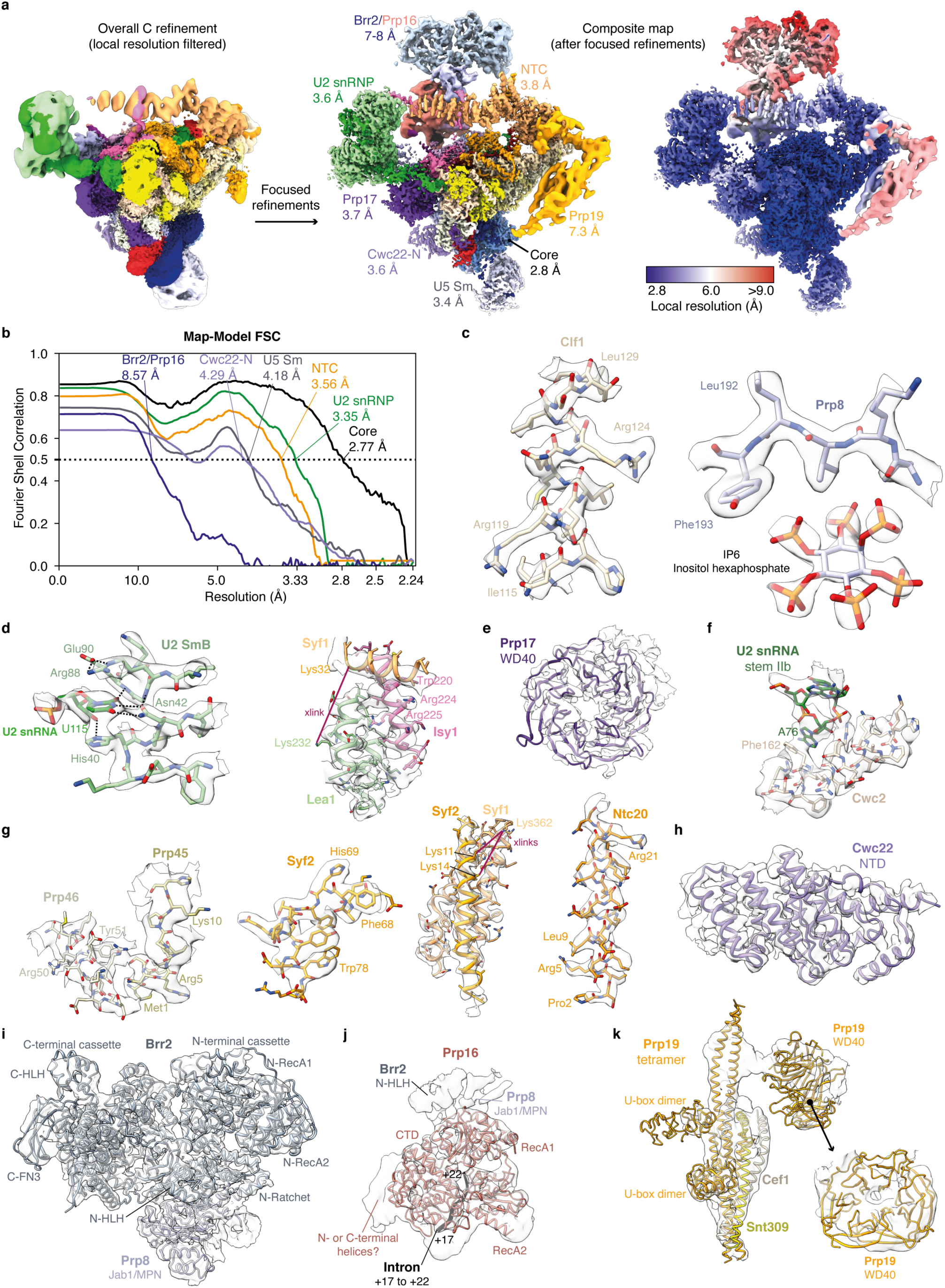
Representative cryo-EM densities for the C complex. **a**, The overall C-complex reconstruction and a composite map of all focussed refinements. Local resolution for each focus-refined map was first calculated in RELION. Individual local-resolution maps were then resampled in Chimera on the overall reconstruction, and at each voxel the local-resolution map with the lowest value (i.e. best resolution) was taken. **b**, FSC curves calculated in PHENIX for each subcomponent of the model (see **Table 1**) against its respective focus-refined map. **c**, Representative densities for proteins and ligands in the C-complex core. Density-modified map is shown. **d-h**, Densities for the focus-refined density-modified maps of the U2 snRNP (**d**), Prp17 WD40 domain, which unambiguously identifies orientation (**e**), Cwc2 (**f**), NTC components (**g**), and for the Cwc22 NTD (**h**). All crosslinks shown are from the B^act^ complex (ref. 73). **i**, Focus-refined Brr2 map. **j**, Focus-refined overall helicase module map around Prp16. **k**, Focus-classified map of the Prp19 tetramer.

**Extended Data Fig. 6.**
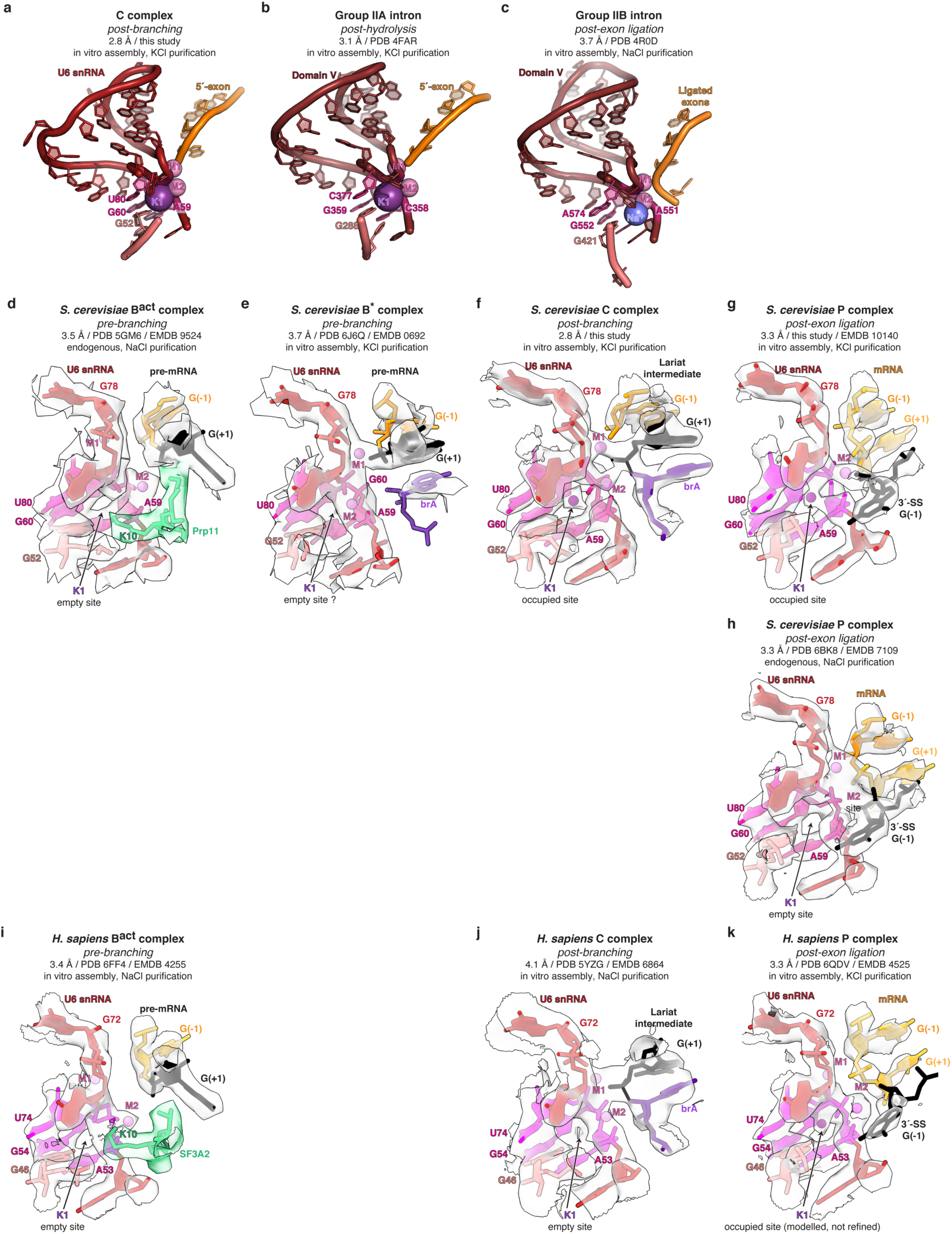
Comparison of key active site elements of the different spliceosomal complexes. **a-c**, Comparison between the proposed M1-M2-K1 C-complex metal cluster and the metal cluster observed in structures of group II introns. A potassium ion stabilises the catalytic centre of the spliceosome. **d-h**, Structure of the M1-M2-K1 catalytic metal cluster in various *S. cerevisiae* spliceosome complexes. The K1 metal cluster is partially blocked in B^act^ by Prp11. Potassium is likely absent at the K1 site in the structure of the P complex assembled endogenously due to the use of NaCl instead of KCl during purification. **i-j**, Structure of the M1-M2-K1 catalytic metal cluster in various *H. sapiens* spliceosome complexes. The K1 metal cluster is partially blocked in B^act^ by SF3A2, the Prp11 homolog. Clear density for potassium is likely not resolved at the K1 site in the structure of the human C complex assembled endogenously due to lower resolution and the use of NaCl instead of KCl during purification. For clarity, in all panels the potassium ion is represented using 0.5x of its Van der Waals radius to allow visualisation of the density. The magnesium ions are represented using their 1x of their van der Waals radii.

**Extended Data Fig. 7.**
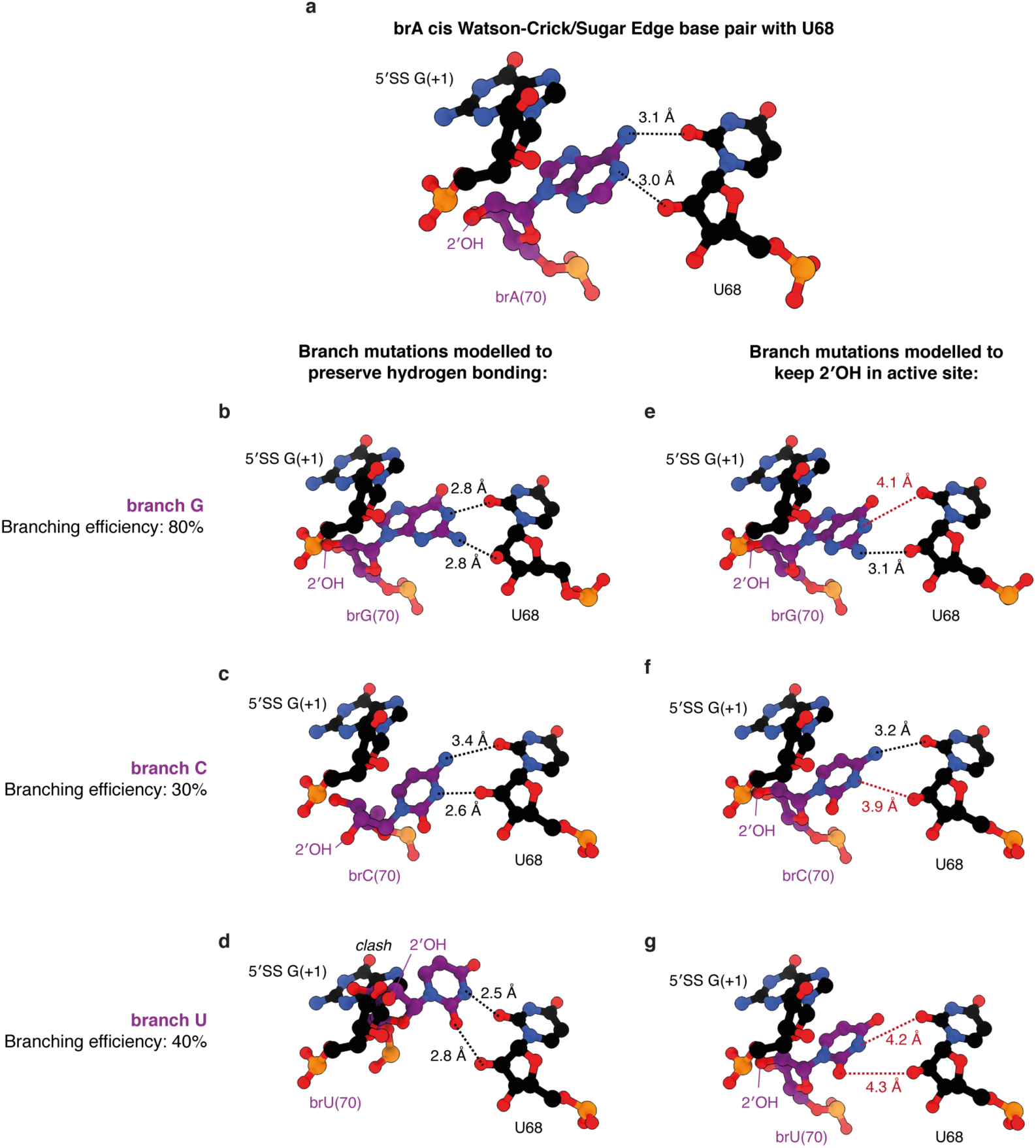
Recognition of brA in C complex. **a**, Proposed model for brA interactions in C complex, as derived from our new high resolution map. **b-g**, Modelling of analogous interactions between mutations at the brA and U68 of the intron. Models **b-d** are based on prototypical examples of analogous base pairs as documented by Leontis and Westhof (ref. 44). Models **e-g** are simple mutations of the branch point base, keeping the sugar and phosphate in the starting position. Branch G can accommodate base pairing and correct 2′OH positioning, while branch C and U can only accommodate either pairing or correct 2′OH positioning.

**Extended Data Fig. 8.**
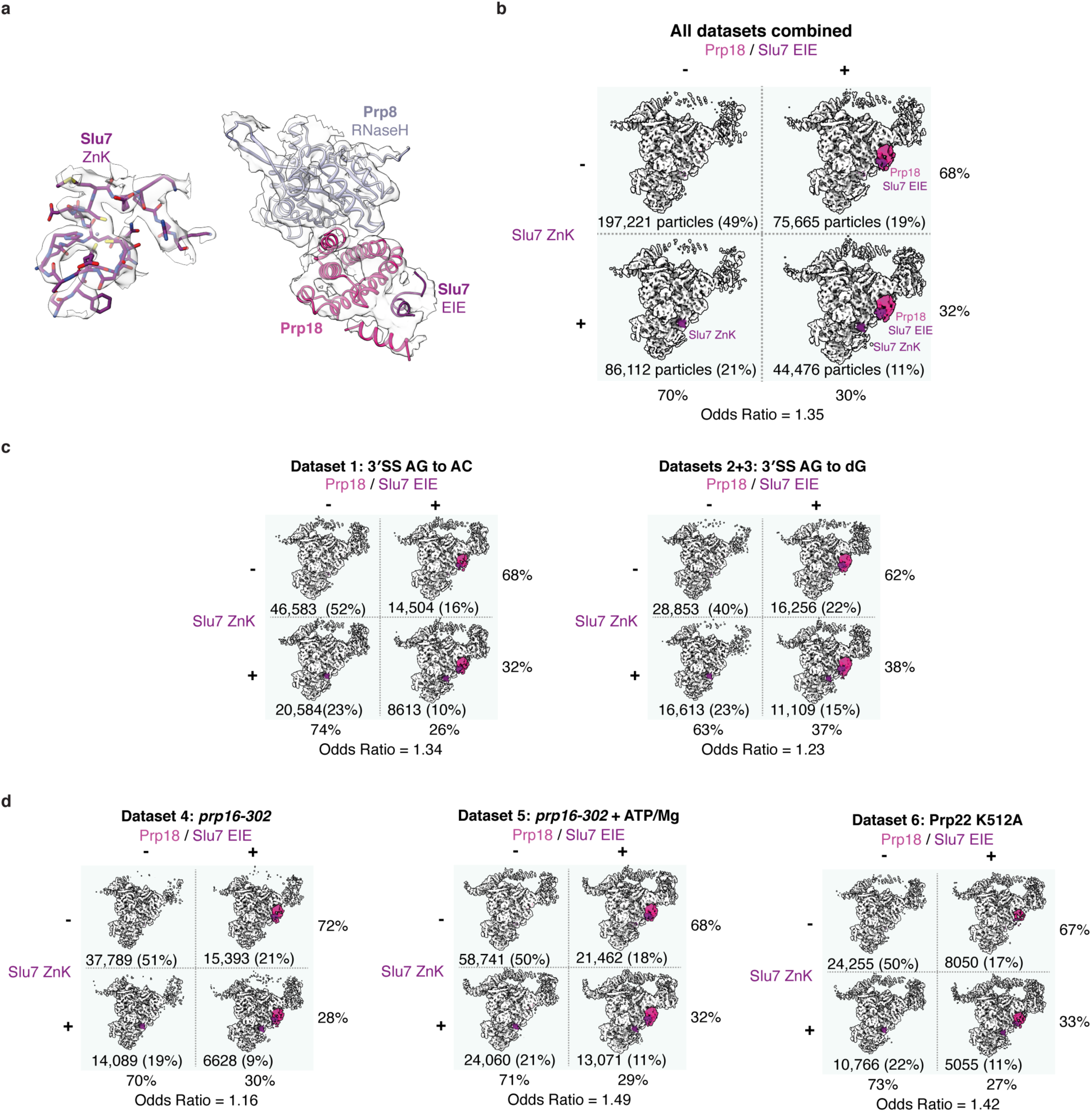
Focussed classification of C-complex particles for the C_i_ conformation. **a**, Density quality for Prp18 and Slu7 ZnK and EIE domains in the final map obtained by classification of all datasets. **b**, Focused classification of Prp18 and the Slu7 ZnK for all combined C-complex datasets. **c**, Focused classification of Prp18 and the Slu7 ZnK for C-complex datasets resulting from impairment of exon ligation by mutation of the 3′-SS. **d**, Focused classification of Prp18 and the Slu7 ZnK for C-complex datasets resulting from impairment of exon ligation by mutation of Prp16 or Prp22.

